# Unsupervised Alignment in Neuroscience: Introducing a Toolbox for Gromov-Wasserstein Optimal Transport

**DOI:** 10.1101/2023.09.15.558038

**Authors:** Masaru Sasaki, Ken Takeda, Kota Abe, Masafumi Oizumi

**Affiliations:** Graduate School of Arts and Science, The University of Tokyo, Meguro-ku, Tokyo, Japan

## Abstract

**Background:** Understanding how sensory stimuli are represented across different brains, species, and artificial neural networks is a critical topic in neuroscience. Traditional methods for comparing these representations typically rely on supervised alignment, which assumes direct correspondence between stimuli representations across brains or models. However, it has limitations when this assumption is not valid, or when validating the assumption itself is the goal of the research.

**New method:** To address the limitations of supervised alignment, we propose an unsupervised alignment method based on Gromov-Wasserstein optimal transport (GWOT). GWOT optimally identifies correspondences between representations by leveraging internal relationships without external labels, revealing intricate structural correspondences such as one-to-one, group-to-group, and shifted mappings.

**Results:** We provide a comprehensive methodological guide and introduce a toolbox called GWTune for using GWOT in neuroscience. Our results show that GWOT can reveal detailed structural distinctions that supervised methods may overlook. We also demonstrate successful unsupervised alignment in key data domains, including behavioral data, neural activity recordings, and artificial neural network models, demonstrating its flexibility and broad applicability.

**Comparison with existing methods:** Unlike traditional supervised alignment methods such as Representational Similarity Analysis, which assume direct correspondence between stimuli, GWOT provides a nuanced approach that can handle different types of structural correspondence, including fine-grained and coarse correspondences. Our method would provide richer insights into the similarity or difference of representations by revealing finer structural differences.

**Conclusion:** We anticipate that our work will significantly broaden the accessibility and application of unsupervised alignment in neuroscience, offering novel perspectives on complex representational structures. By providing a user-friendly toolbox and a detailed tutorial, we aim to facilitate the adoption of unsupervised alignment techniques, enabling researchers to achieve a deeper understanding of cross-brain and cross-species representation analysis.

## Introduction

A central topic in neuroscience is the comparison of how sensory stimuli are represented in different brains [1, 2]. These representations are typically assessed through multi-dimensional neural activity patterns elicited by sensory stimuli, derived from neural recordings. In addition to neural activity data, behavioral data from psychological experiments are also used to estimate these representations. Common comparisons include those between different individuals within the same species, between different species, or even between biological brains and artificial neural network models. Such studies help to uncover similarities or differences in how humans, other animals, or artificial neural networks represent sensory stimuli, and ultimately contribute to our understanding of the computational principles underlying these representations.

Traditionally, the comparison between different representations has been performed in a supervised manner, using methods known as supervised alignment.

Representational similarity analysis (RSA) [1, 3] is one of the most widely used methods in neuroscience that falls into this category. Supervised methods compare neural representations based on the assumption that the representations of stimuli in one brain directly correspond to the representations of the same stimuli in the other brain (Fig. 1(a)). For example, the representation of an apple in one brain is assumed to correspond directly to the representation of an apple in the other brain. Under this assumption, the representations of various sensory stimuli are quantified by correlations between representational similarity matrices [3], whose elements represent the similarity between representations, or by the Procrustes alignment [4] between representational structures.

**Fig 1.**
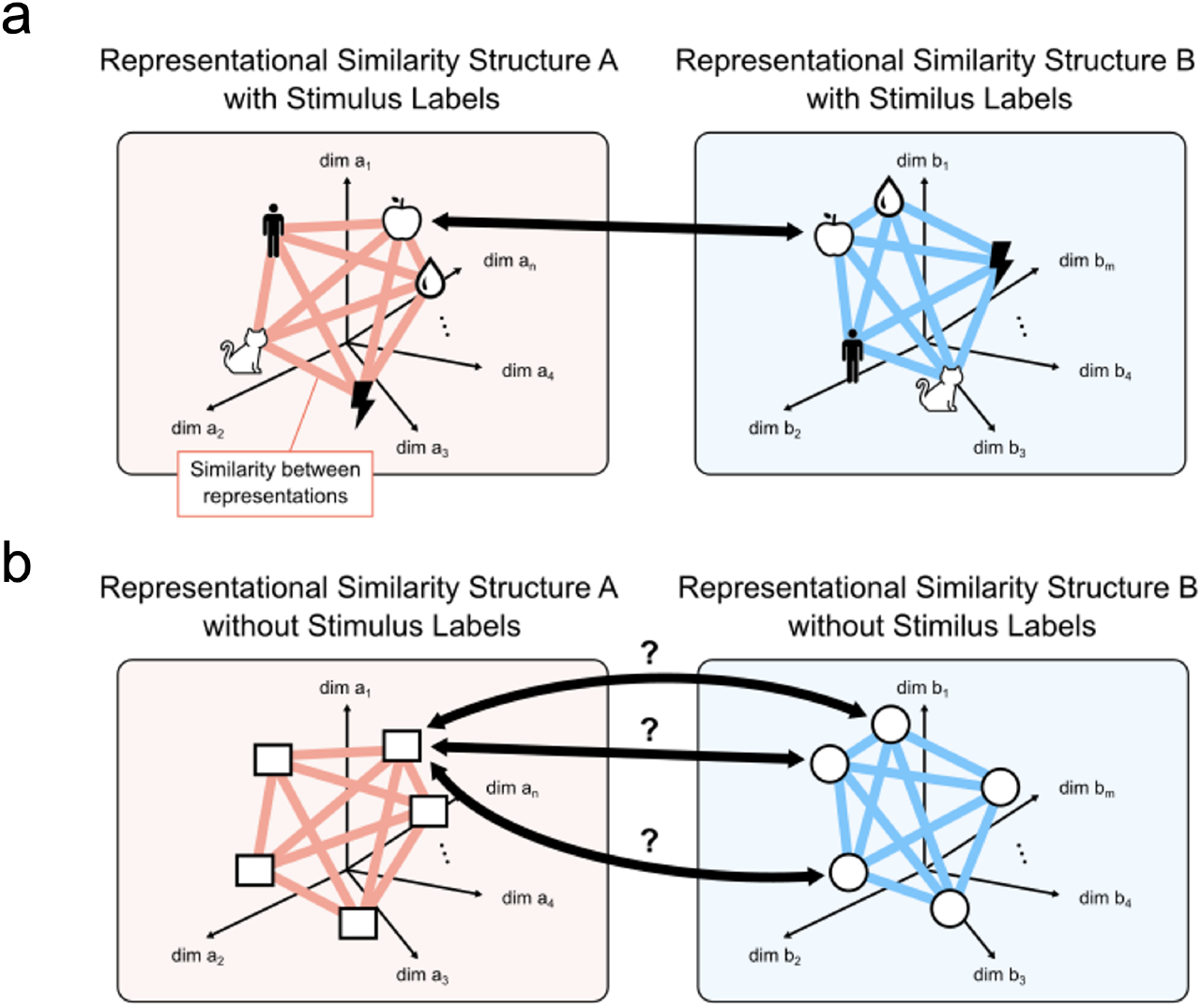
Comparing different representations. (a) Supervised alignment: Assuming correspondence between stimulus labels, different representational similarity structures are compared and aligned to assess the similarity between the representational similarity structures. (b) Unsupervised alignment: Different representational similarity structures are compared and aligned without the use of stimulus labels, relying solely on each internal similarity structure. The optimal correspondence between stimuli is found through the optimization process.

Although supervised alignment provides a simple and useful measure of overall similarity between different representations, it has limitations when the assumption of direct correspondence between identical stimuli is not valid, or when validating this assumption itself is the goal of the research. For example, in our previous work [5], we explored the possibility that behavioral representations of color stimuli, estimated from color similarity judgment data, could differ between individuals, a concept that parallels the “inverted qualia” thought experiment in consciousness research. Although the assumption of direct correspondence may be reasonable to some extent within similar populations (e.g., color-neurotypical human participants), it cannot be guaranteed across populations with different characteristics (e.g., between color-neurotypical and color-atypical participants) or across species (e.g., between humans and other animals). Even when comparing apparently similar representations, validation of the assumption itself is valuable, as it can serve as evidence of strong structural commonalities between the representations.

To overcome the limitations of supervised comparison, we consider that unsupervised alignment methods should be used for a wide range of applications in neuroscience. Unsupervised alignment does not assume predefined correspondences between the representations of identical stimuli, but instead attempts to find optimal correspondences through an optimization process (Fig. 1(b)). This approach can distinguish between different types of structural correspondences that supervised methods like RSA cannot, such as one-to-one mapping, coarse group-to-group mapping, or consistently shifted one-to-one mapping (see the next section for details).

This paper provides a methodological framework for unsupervised alignment in neuroscience using Gromov-Wasserstein Optimal Transport (GWOT), aiming to facilitate its application in diverse neuroscience scenarios. The GWOT, proposed in [6], is based on the mathematical theory of optimal transport [7], which finds the correspondence between the point clouds in different domains based only on the internal relations (distances) between the points in each domain and thus does not rely on external labels. The GWOT method has been applied with great success in various fields, including the alignment of 3D objects [6], the translation of vocabularies in different languages [8, 9], and the matching of single cells in single-cell multi-omics data [10]. On the other hand, in neuroscience, to the best of our knowledge, we are the first to provide a framework of unsupervised alignment with GWOT for comparing neural representations. Although GWOT has been used for the purpose of aligning brain shapes between individuals [11], there was no previous work that used it for the purpose of aligning neural representations. By using the present toolbox, previous studies from our group reported the unsupervised alignment of behavioral similarity structures of color [5, 12], natural objects [13], and electrophysiological recordings data in mice for natural scenes and natural movie stimuli [14]. In the present paper, in addition to applying it to several neuroscience-related datasets, we also conduct a simulation to demonstrate the theoretical differences between GWOT and conventional RSA, a method widely used in neuroscience. The simulations are helpful in understanding not only the methodology, but also the significance of the GWOT in comparison to the conventional method.

To effectively demonstrate the framework of unsupervised alignment based on GWOT, this paper introduces an end-to-end tutorial along with a user-friendly toolbox, named GWTune, which is a crucial resource for making this unsupervised alignment method accessible to a wider field. The tutorial provides clear, step-by-step guidance, covering the entire process from data preparation to optimization and evaluation, enabling readers to easily grasp the key concepts and procedures at each stage. Moreover, all functionalities are seamlessly integrated into the toolbox, ensuring that even researchers with limited programming experience can effectively utilize it for their analyses.

GWTune provides more flexibility for general applications by using random initial value search and effective hyperparameter tuning. One of the challenges of using GWOT is that it involves non-convex optimization and requires careful hyperparameter tuning. To address this, we introduced random initialization of the transportation matrix, which helps avoid local minima and increases the likelihood of finding the global minimum, thereby improving the optimization process. Also, our toolbox supports user-friendly hyperparameter tuning by integrating Optuna [15], an advanced hyperparameter optimization tool. Using random initialization and Optuna for hyperparameter tuning, our approach enhances the overall robustness and efficiency of the optimization process—an integration that is unique and not found in previous studies [10, 16]

To demonstrate the utility of GWOT-based unsupervised alignment and our toolbox, we present applications in three key data domains in neuroscience: behavioral data, neural data, and neural network models. For behavioral data, we analyzed psychological similarity judgments of natural objects from the THINGS dataset [17–19], using GWOT to align similarity structures between male and female participant groups. For neural data, we applied GWOT to align similarity structures of neural responses to visual images from mouse Neuropixels recordings provided by the Allen Brain Institute [20, 21]. In the domain of neural network models, we used GWOT to align internal representations of visual deep neural networks (DNNs), specifically between ResNet50 [22] and VGG19 [23].

We expect that this tutorial and the accompanying toolbox will provide a robust solution for various research applications in neuroscience and other fields, making the powerful GWOT methodology accessible to a wider range of users. The open source code and example data for the toolbox are available on GitHub at https://oizumi-lab.github.io/GWTune/.

## Alignment in neuroscience

In this paper, we primarily consider the problems of alignment in neuroscience (Fig. 2), but the toolbox is readily applicable to data in other research areas. Data in neuroscience for which alignment is considered in the current literature can be categorized into three key main targets: neural data, behavioral data, and artificial neural network models (Fig. 2(a)). Neural data include any kind of recordings of neural activity, such as Neuropixels, Ca imaging, EEG, ECoG, fMRI, etc. Behavioral data include any kind of behavioral reports, e.g. similarity judgments and ratings, multiple choice, free verbal or written descriptions, etc. Neural network models include any type of neural network model, e.g. convolutional neural networks, recurrent neural networks, transformers, etc. There may be other types of data to which the alignment framework in this paper is applicable, but here we consider these three main targets for simplicity and clarity.

**Fig 2.**
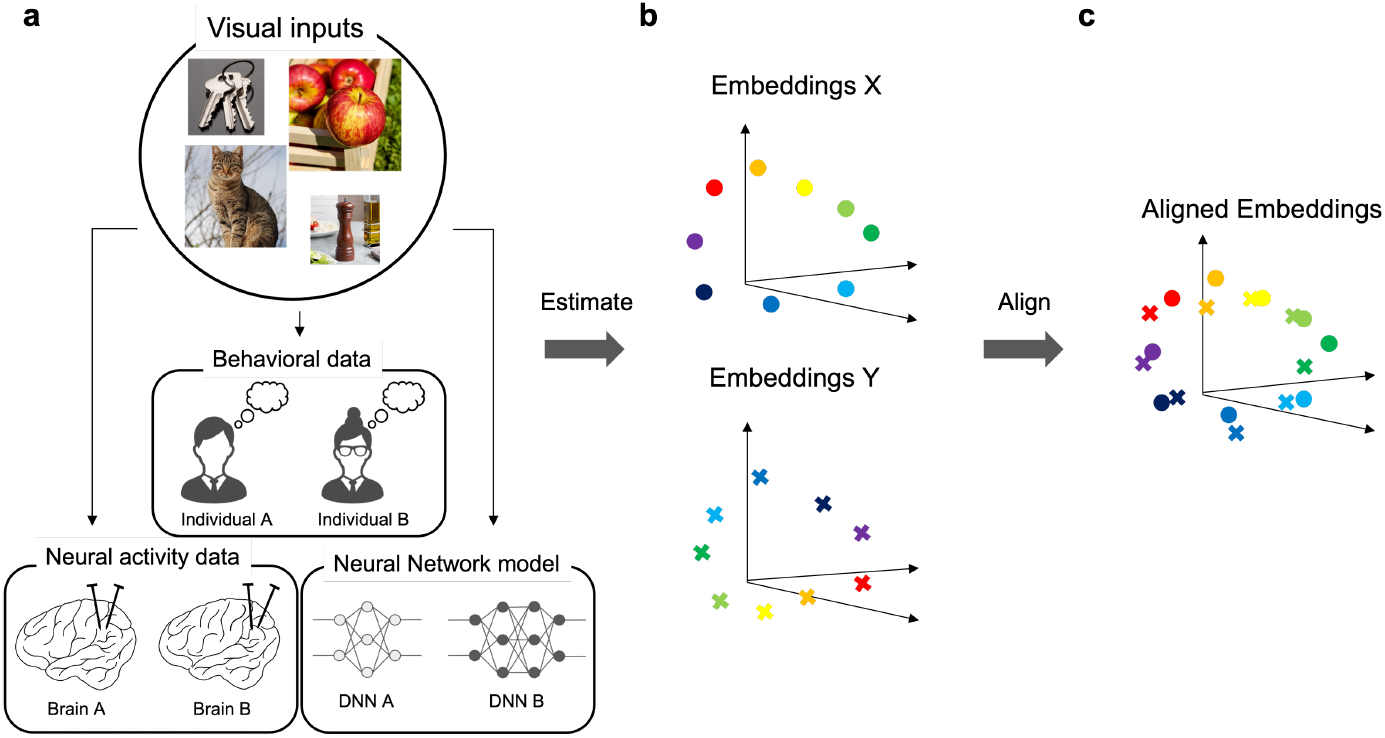
Alignment in neuroscience. (a) Three data domains in neuroscience. Behavioral data, neural activity data, and neural network models. As examples, behavioral responses in different individuals, neural responses in different brains, and model neuron activities in different deep neural network (DNN) models to visual inputs are considered. (b) Embeddings estimated from the three data domains. (c) Alignment of the two embeddings.

Despite the variety of data types, we consider the same common alignment problem in this paper: the alignment of point clouds or “embeddings” (Fig. 2(b)). A point cloud is a set of data points in space, usually defined by a set of coordinates in space. In this paper, we use the common terminology “embeddings” to mean points in space, regardless of whether the data come from neural recordings, behavioral experiments, or neural network models. To perform an alignment, we should first convert raw data into a set of points in space. For example, in the case of neural data or neural network models, we can simply use the firing rates of real neurons or the activities of model neurons to a set of different stimuli as points in space, where the dimension represents the activity of each neuron. In the case of behavioral data, we typically estimate so-called psychological embeddings [18, 24] that best explain the behavioral responses to different stimuli. After obtaining the sets of estimated embeddings, we align the embeddings, typically by minimizing some objective function, such as the L2 distance between the embeddings after the alignment (see Supporting Information for details).

### Supervised alignment and unsupervised alignment

There are two broad categories of alignment methods: supervised alignment and unsupervised alignment. Both are techniques for aligning two sets of embeddings, but they differ in whether or not they assume correspondences between the embeddings of the different sets. The former uses predefined correspondences to perform the alignment, while the latter does not rely on any correspondences.

#### Supervised alignment

A typical and simplest supervised alignment method is the Procrustes analysis [25]. This simply finds the best rotation matrix that minimizes the L2 distance between two different sets of embeddings, given the correspondences between the embeddings across the different sets (see the Supporting Information for details). The minimized L2 distance represents the similarity between the sets of the embeddings.

There is another most widely used method to compare and quantify the similarity between the sets of embeddings in neuroscience, called Representational Similarity Analysis (RSA) [3, 26, 27]. Although RSA is technically not an alignment method in the sense that it does not explicitly align the embeddings, we could classify this method in the same category as supervised alignment, or more generally, we can call it a supervised comparison method. RSA directly compares dissimilarity matrices of two sets of embeddings by evaluating how similar they are. The elements of the dissimilarity matrix *D* are represented as *D*_*ij*_ *= f*_dis_ (***x***_*i*_, ***x***_*j*_), where *f*_dis_ is a function that quantifies the dissimilarity between embeddings *i* and *j*, such as the Euclidean distance or the cosine distance. Then, based on two dissimilarity matrices *D* and *D*′, the dissimilarity between them is evaluated by a function *g*_dis_ (*D, D*′). In this paper, we use a simple RSA method, where *g*_dis_ is Pearson’s correlation coefficient, to compare dissimilarity matrices. See [28] for a comprehensive list of dissimilarity metrics categorized in supervised alignment and a mathematical characterization of the metrics.

#### Unsupervised alignment

In contrast, the methods of unsupervised alignment compares two sets of embeddings without given correspondences, but instead, find the correspondences during the optimization process. Although unsupervised alignment methods have not been yet applied in many cases in neuroscience (e.g. [5, 11–14]), they will potentially find various applications. As we will show later in Results section, unsupervised alignment methods can reveal more detailed structural similarities or differences than supervised alignment.

With unsupervised alignment, which finds the optimal correspondences without using any labels, there are several possible scenarios that can never be revealed by supervised alignment: In principle, even if there are equally high correlations between the similarity structures of two sets of embeddings in terms of supervised comparison, the following qualitatively different scenarios are possible, as shown in Fig. 3. (a) Same one-to-one mapping between one set of embeddings *X* and the other set of embeddings *Y* (Fig. 3(a)): The embeddings of the same stimuli correspond to each other one-to-one (e.g., the embedding of an apple in *X* corresponds to that of an apple in *Y*). This indicates that the representations are relationally “equivalent” across the different sets of embeddings *X* and *Y*. (b) Partially different one-to-one mapping (Fig. 3(b)): The embeddings of some stimuli correspond to each other one-to-one, but not all. For example, the embedding of an apple in *X* corresponds to the embedding of an orange in *Y*, while other embeddings, such as a cat in *X*, still correspond to a cat in *Y*. This scenario indicates that while there is a generally consistent mapping, certain stimuli have non-identical correspondences between the two sets of embeddings. (c) Coarse group-to-group mapping (Fig. 3(c)): The embeddings correspond to each other at the group level, but not at the fine-item level. For example, the embeddings of fruits in *X* correspond to those of fruits in *Y*, but the embedding of an apple in *X* does not correspond specifically to an apple in *Y*. Instead, it corresponds to any fruit in *Y*, indicating a more generalized correspondence at the category level rather than at the individual item level. (d) Consistently shifted one-to-one mapping (Fig. 3(d)): Most embeddings correspond to each other one-to-one, but are systematically shifted. For example, the embedding of an apple in *X* consistently corresponds to the embedding of an orange in *Y*, and similarly, a cat in *X* consistently corresponds to a dog in *Y*. This scenario suggests a consistent but shifted mapping pattern, where each stimulus has a consistent but different counterpart in the other set. Shifted mapping would occur, for example, when the embeddings have a low-dimensional symmetric structure, such as a circle. As we will show later in a concrete simulation, supervised comparisons such as RSA cannot distinguish these possible cases.

**Fig 3.**
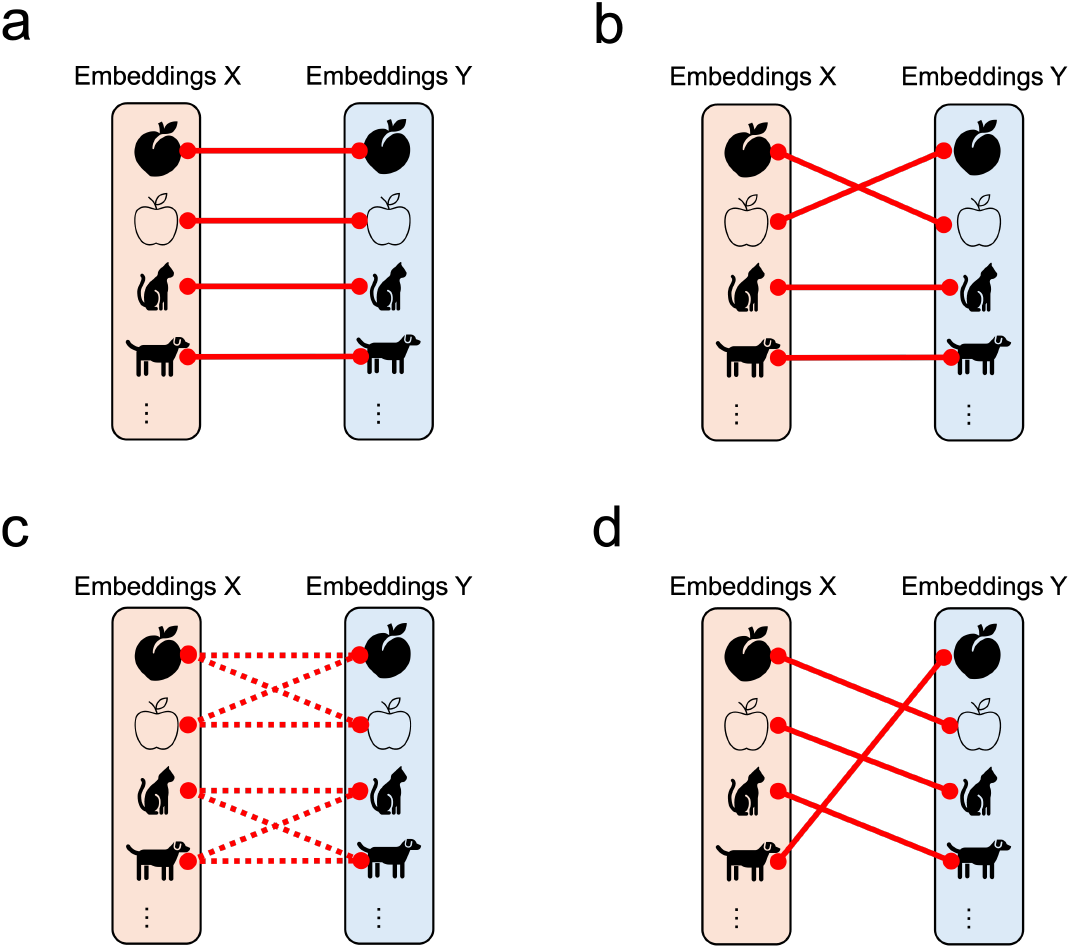
Possible consequences of unsupervised alignment. (1) Same one-to-one mapping. (2) Partially different one-to-one mapping. (3) Coarse group-to-group mapping. (4) Shifted one-to-one mapping.

## Gromov-Wasserstein Optimal Transport (GWOT)

Gromov-Wasserstein optimal transport [6] is an unsupervised alignment technique that finds correspondence between two point clouds (embeddings) in different domains based only on internal distances within the same domain. Mathematically, the goal of the Gromov-Wasserstein optimal transport problem is to find the optimal transportation plan Γ between the embeddings in different domains, given two dissimilarity matrices *D* and *D*′within the same domains (Fig. 4). The transportation cost, i.e., the objective function, to optimize in GWOT is given by

**Fig 4.**
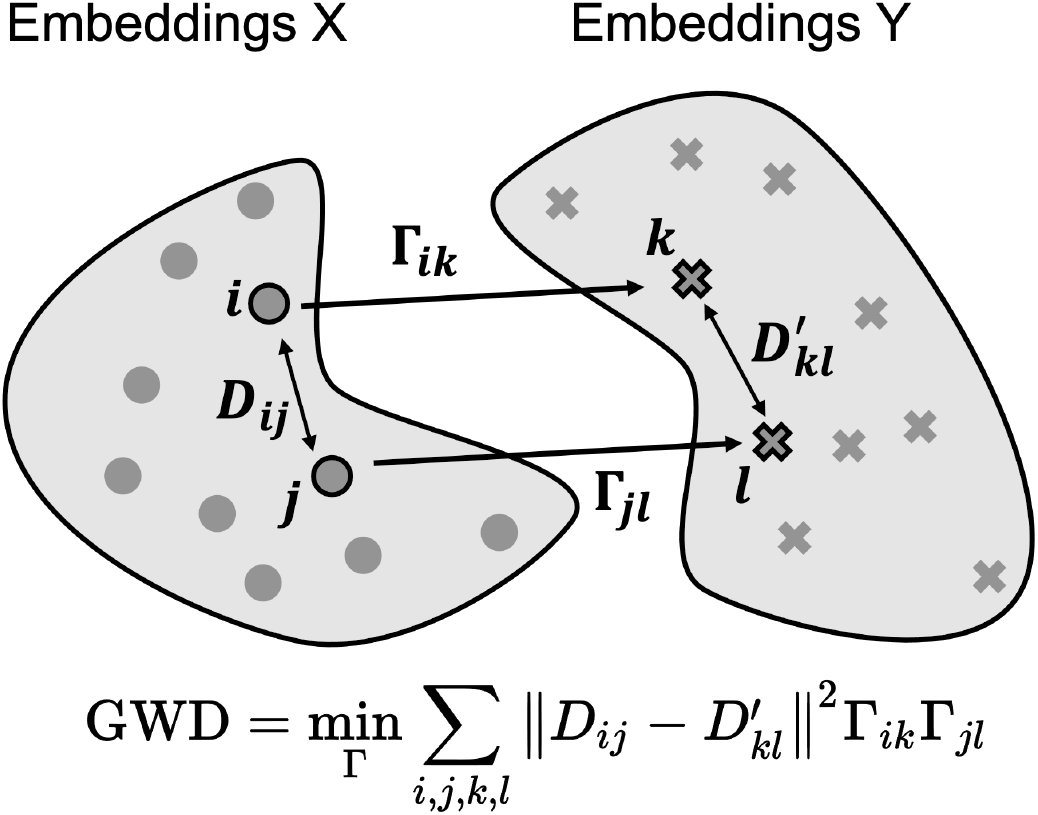
Unsupervised alignment using GWOT. Schematic of Gromov-Wasserstein optimal transportation (GWOT). *D*_*ij*_ represents the distance between the *i*th embedding and the *j*th embedding in one set of embeddings *X* and 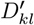 represents the distance between the *k*th embedding and the *l*th embedding in the other set of embeddings *Y*. Γ_*ik*_ represents the amount of transportation from the *i*th embedding in *X* and the *k*th embedding in *Y*.

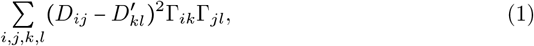

where a transportation plan Γ must satisfy the following constraints: ∑_*j*_ = Γ_*ij*_ *= p*_*i*_, ∑_*j*_ = Γ_*ij*_ *= q*_*j*_, and ∑_*j*_ = Γ_*ij*_ *=* 1, where ***p*** and ***q*** is the source and target distribution of mass for the transportation problem, respectively, whose sum is 1. Under this constraint, the matrix Γ is considered as a joint probability distribution with the marginal distributions being ***p*** and ***q***. As for the distributions ***p*** and ***q***, we set ***p*** and ***q*** to be the uniform distributions. Each entry Γ_*ij*_ describes how much of the mass on the *i*-th point in the source domain is transported onto the *j*-th point in the target domain. The entries of the normalized row 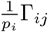 can be interpreted as the probabilities that the embedding ***x***_*i*_ corresponds to the embeddings ***y***_*j*_.

Gromov-Wasserstein distance (GWD) is defined as the minimized transportation cost (Eq. 1 with respect to transportation plans Γ:

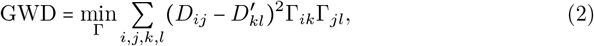

In short, due to the term 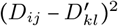 in Eq. 2, the objective function is minimized when the transportation plan Γ aligns the point clouds in the two domains such that the distances between the points in one domain are similar to the distances between the corresponding points in the other domain. Although we may apply the original GWOT to some applications, in this paper, we mainly consider a more general version of GWOT, called entropic GWOT, which includes an entropy-regularization term in the objective function.

### Entropic GWOT

Previously, it was shown that adding an entropy regularization term can improve computational efficiency and help identify good local optimums of the Gromov-Wasserstein optimal transport problem [7, 29]. This entropic GWOT is formulated as

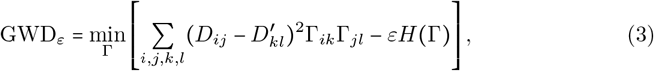

where *H (*Γ) is the entropy of the transport plan Γ. The original GWOT is a special case of Eq. 3 when *ε* = 0. The hyperparameter *ε* controls the tradeoff between the cost of transportation and the entropy of the transportation plan. A larger value of *ε* makes the solution smoother (denser), while a smaller value of *ε* provides a solution closer to the original problem (Eq. 2), but makes the solution sharper (sparser). Choosing appropriate epsilon values helps to find good local optimums while avoiding getting stuck in bad local optimums.

#### Hyperparameter tuning of ε and initialization of transportation plan Γ

To find good local optimums, we need to perform hyperparameter tuning on *ε* in Eq. 3 to adjust the trade-off between small and large epsilon values. After many iterations of hyperparameter search, we would obtain many local optimal solutions for the transportation plan. Among the local optimums, we select the transportation plan that minimizes the Gromov-Wasserstein distance without the entropy regularization term (Eq.2) as the global optimal solution, following the procedure proposed in a previous study [10].

In some applications, it may not be sufficient to simply search for the hyperparameter *ε* with the fixed initialization of the transport plan Γ, but it is also necessary to search for different initializations. To avoid getting stuck in bad local minima, it is effective to try many random initializations of transportation plans. In our toolbox, we recommend searching for good local minima by simultaneously changing both *ε* and the initialization of the transport plan Γ as the default option. See Supporting Information for details on hyperparameter tuning on *ε* and initialization.

## Comprehensive guide to entropic GWOT

### Process overview

The main aim of this paper is to provide a comprehensive tutorial on unsupervised alignment using entropic GWOT for the neuroscience community and related fields. For this purpose, we also provide a user-friendly toolbox for entropic GWOT to enable effective and user-friendly hyperparameter tuning. Our toolbox uses two key Python libraries: Optuna [15] for hyperparameter tuning and POT (Python Optimal Transport) [30] for entropic GWOT computation. Our toolbox, including all example data used in the paper, is available at https://oizumi-lab.github.io/GWTune/).

The whole process of unsupervised alignment using GWOT is summarized in Fig. 5.

**Fig 5.**
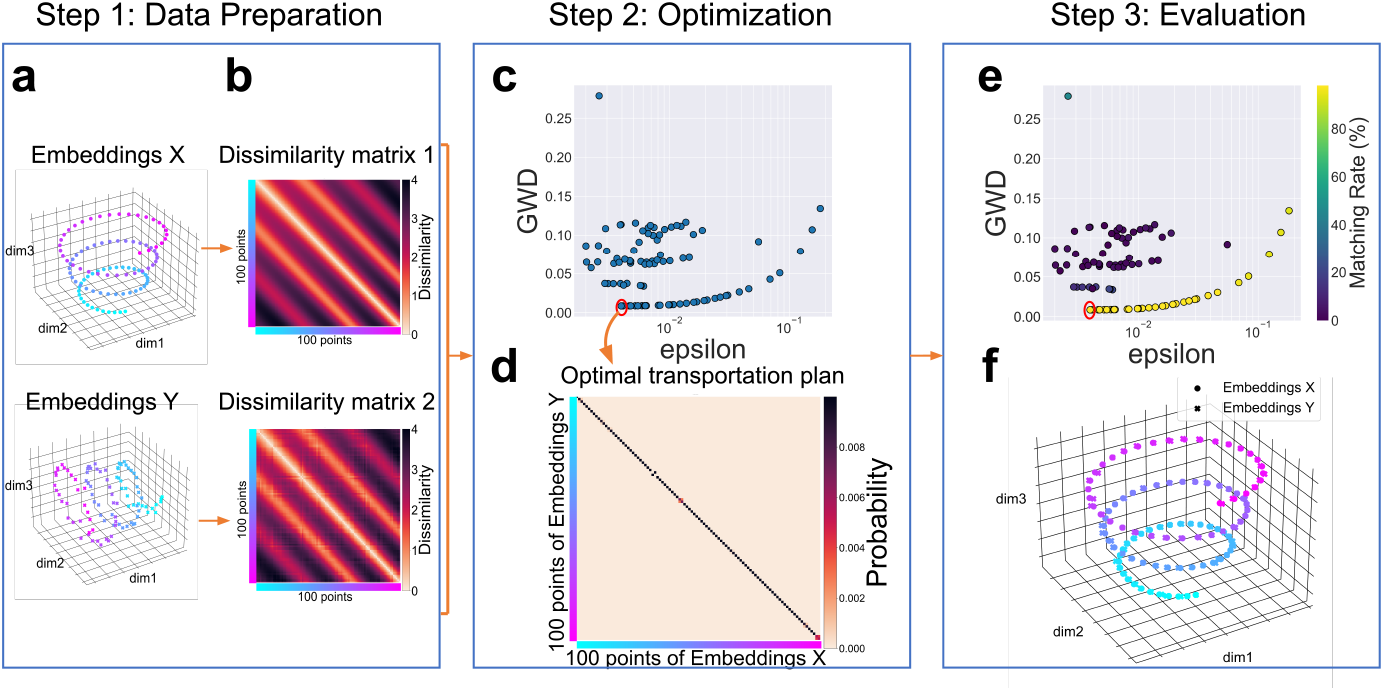
Complete workflow for unsupervised alignment with entropic GWOT. (a) Embeddings of *X* and *Y*. (b) Dissimilarity matrices of embeddings *X* and *Y*. (c) Optimization results showing the relationship between GWD and the hyperparameter *ε*. (d) Optimal transportation plan Γ with the lowest GWD value, selected from the optimization results shown in the red circle in (c) and (e). (e) Optimization results showing the relationship between the epsilon values and GWD values (the same as (c)) with the matching rate of its transportation plan using the colorbar in (e). The matching rate is the the frequency of the diagonal elements that are the highest among the other elements with the same rows. (f) Visualization of the embeddings X and the aligned embeddings Y in 3D space.

1. The dissimilarity matrices are computed from the embeddings.
2. Optimization of GWOT is performed between the dissimilarity matrices.
3. The embeddings are aligned by using the optimal transportation plan Γ found by GWOT.

Below, we provide a detailed explanation of each step and pseudo code to perform each step based on our toolbox.

#### Step 1: Preparation of dissimilarity matrices

As Step 1, dissimilarity matrices should be prepared for use in the subsequent unsupervised alignment. For some experimental data, dissimilarity matrices may be directly available; however, in other cases, only embeddings, such as neural activity data, are available. In these cases, the dissimilarity matrices need to be computed based on the distances between the embeddings. There are many ways to compute the distances, including cosine distance, Euclidean distance, and inner product. See the specific examples of embeddings and dissimilarity matrices in the “Practical applications and case studies” section.

We demonstrate Step 1 using synthetic data. We generated 100 points on a spiral in 3D space and considered them as “Embeddings *X*” (Fig. 5(a)). Then, we added some independent noise to the three dimensions of all points in the Embeddings *X*, rotated them by 90 degrees around the axis of dim 2, and considered them as “Embeddings *Y* “(see Fig. 5(a)). The points with the same colors in *X* and *Y* in Fig. 5(a) indicate the same points of the spiral. Based on these embeddings, the dissimilarity matrices are computed by the Euclidean distance of the embeddings *X* and *Y*. The two dissimilarity matrices are visualized in Fig. 5(b) where the colorbar shows the dissimilarity values (the Euclidean distance) for each pair of embeddings, and the colors next to the two axes in Fig. 5(b) represent the points with the same colors shown in Fig. 5(a). These dissimilarity matrices are used as input for the optimization of GWOT in Step 2.

#### Step 2: Optimization and hyperparameter tuning

As Step 2, the hyperparameter *ε* values and initial transportation plans are searched for the optimization of GWOT. The *ε* values act as regularization parameters in Eq. 3, where larger values makes the solution smoother (denser) and smaller values makes the solution sharper (sparser). We search for the optimized hyperparameter *ε* values from a certain range of *ε* values. Additionally, the choice of the initial transportation plan is crucial, as it can lead to different local minima. Exploring a wide variety of initial transportation plans increases the chances of avoiding poor local minima and finding better global optima. A standard way we recommend to initialize a transportation plan is to generate it randomly. However, for some problems, one can also consider initializing transportation plans using other prior knowledge. See the Supporting Information for details on hyperparameter tuning for *ε* and initializing transportation plans.

We demonstrate Step 2 using the two dissimilarity matrices created in Step 1. We set the range of *ε* from 0.002 to 0.2 and the number of the *ε* samples to 100. As for the sampling of *ε*, we used the Tree-Structured Parzen Estimator, an effective Bayesian sampling method. As for the initialization of the transportation plans, we randomly generated initial transportation plans for each *ε* value. After 100 iterations of *ε*, we obtained the optimization results shown in Fig. 5(c). Each dot in Fig. 5(c) shows the GWD values computed from the optimal transportation plans found for the sampled *ε* values. Among the optimal transportation plans, i.e., local optimums, found in the iterations, we choose the one with the minimum GWD, shown in the red circle in Fig. 5(c), as the global optimum. The optimal transportation plan with the minimum GWD was found around epsilon values of 4.0 × 10 ^−3^. Fig. 5(d) shows the optimal transportation plan with the minimum GWD in Fig. 5(c) between the dissimilarity matrices of Embeddings *X* and Embeddings *Y*. The colorbars next to the two axes in Fig. 5(d) indicate the colors of the data points in the two embeddings in Fig. 5(a).

#### Step 3: Evaluation and visualization of the optimization results

As Step 3 (the last step), the optimization results should be evaluated from several perspectives. Specifically, we provide the following evaluation methods: matching rate of the optimal transportation plan, inspection of other local minima, and visualization of aligned embeddings. The matching rate of the optimal transportation plan is computed by comparing the empirically found correspondence, represented by the elements of the optimal transportation plan, with a known ground truth or correspondence. If the known correspondences are categorical, this is referred to as the “categorical matching rate”. Inspecting other local minima helps to understand the landscape of the optimization problem and to evaluate the robustness of the solutions. Visualization of aligned embeddings involves plotting the embeddings in 2D or 3D space after alignment. Using dimensionality reduction techniques such as PCA, the embeddings can be projected into a lower dimensional space, allowing visual inspection to ensure that corresponding embeddings are close together, which helps to validate the quality of the alignment. See the “Methods” section of Supporting Information for details.

We demonstrate Step 3 by using the optimization results obtained in Step 2. First, by observing the optimal transportation plan with the minimum GWD in Fig. 5(d), we can see that most of the diagonal elements have high values, indicating that most of the points in *X* correspond to the same points in *Y* with high probability. This result means that the two similarity structures are almost perfectly matched at the level of individual points by GWOT. The top-1 matching rate, which is the frequency of the diagonal elements that are the highest among the other elements in the same rows, is 98%. We can evaluate the robustness of this optimal solution by examining other local minima. Fig. 5(e), which is the same as Fig. 5(c), adds the information about the top-1 matching rate of the other local minima with colors. This figure suggests that the optimal transportation plans with high matching rate can be found robustly, because the GWD values of the optimal transportation plans with high matching rate (points with blueish colors) are far enough away from the GWD values with low top-1 matching rate (points with yellowish colors). Finally, we visualize the quality of unsupervised alignment based on GWOT by plotting Embeddings *X* and Embeddings *Y* in the same space (Fig. 5(f)). Fig. 5(f) shows that every data point with the same color in the two embeddings is located close to each other. This result indicates that the two embeddings are well aligned by the unsupervised alignment based on GWOT.

#### Pseudo Code

We provide a Python pseudo code that summarizes the process from Step 1 to Step 3 in Algorithm 1. This code is pseudo code in the sense that it cannot be executed as is, but the names of the classes or methods are real and are implemented in our toolbox. As Step 1, two data of embeddings are stored in the class `Representation`. Next, as Step 2, some important parameters such as `sampler` (the sampler method), `init_matrix` (the initialization method), `num_trial` (the number of hyperparameter tuning trials on *ε*), and `epsilon_list` (the range of *ε*) are set by using the class `OptimizationConfig`. Here, for example, the sampler method is set to `tpe`, which is Tree-Structured Parzen Estimator, and the initialization method is set to `random`, which is random initialization. Then, the GWD is optimized by applying the `gw_alignment` method to the `AlignRepresentations` class object. As a result, for each *ε* value, the optimized transportation plans and the corresponding GWDs are obtained. For Step 3, the matching rate is computed by calc_accuracy method, and the inspection of other local minima and visualization of aligned embeddings are done by `show_optimization_log` and `visualize_embedding` methods, respectively.

##### Algorithm 1 Pseudo code of optimization of GWOT

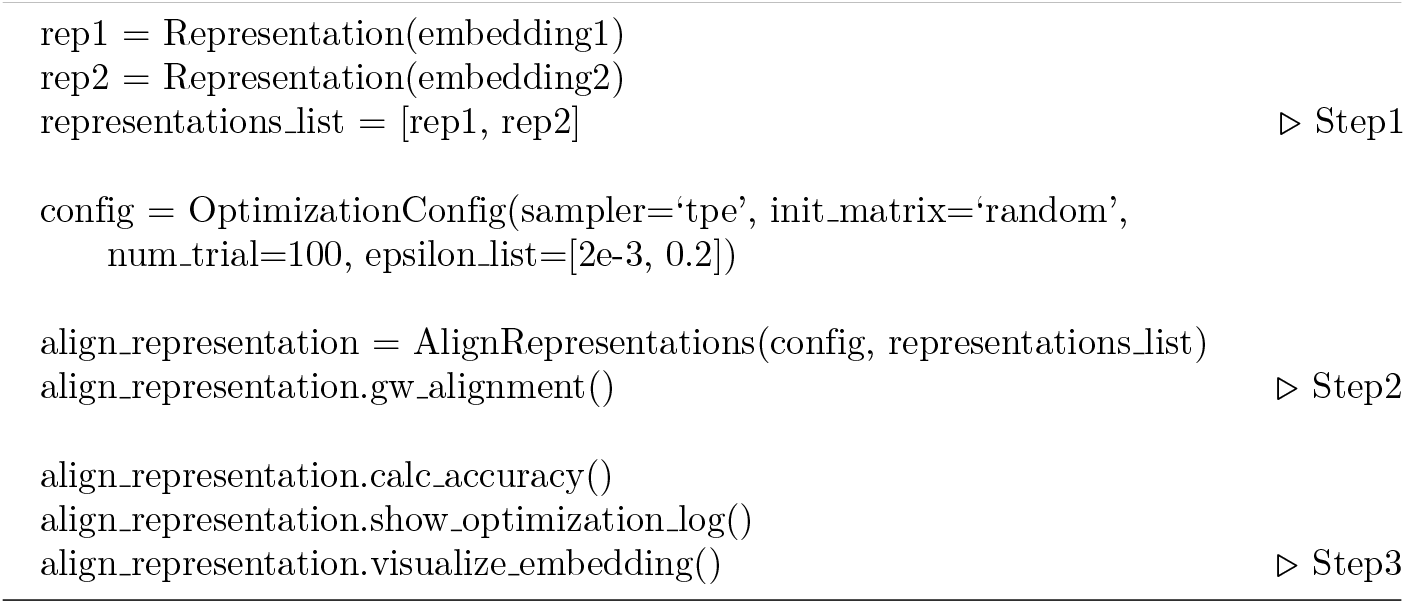

## Practical applications and case studies

Here, as a tutorial, we will demonstrate the application of our toolbox to three key domains of data in neuroscience: behavioral data, neural data, and models. Before analyzing real data, we will first illustrate two important concepts using synthetic data: 1) the impact of the initialization strategies and 2) the difference between unsupervised alignment and supervised alignment. We will then demonstrate the analysis on real data (see Supporting Information for a comparison of the initialization strategies).

### Synthetic data illustrating the impact of the initialization strategies

In this toolbox, we have introduced the random initialization for the transportation matrix. The primary advantage of using random initialization over a fixed uniform distribution is its ability to explore a broader range of possible solutions, thereby increasing the likelihood of finding a better local or even global optimum. Since the optimization process of GWOT is inherently non-convex, there are many opportunities for it to get stuck in suboptimal local minima. Introducing variability in the initial values can help overcome this issue by enabling the optimization process to explore a wider solution space.

To explore the impact of initialization strategies on optimization performance, we conducted a controlled simulation. The purpose of this experiment was to create a scenario with multiple local minima by constructing symmetric shapes, which could cause the optimization process of GWOT to get stuck in suboptimal solutions. We aimed to investigate how different initialization strategies could affect the likelihood of finding a better solution, specifically the global optimum, in such situations.

In this experiment, we generated synthetic data by creating a circle with asymmetric perturbations (Fig. 6a) and added noise to create a second circle (Fig. 6b). Each point was assigned an index. Due to the high symmetry of the circles, GWOT is prone to getting stuck in non-optimal local minima, which can correspond to rotated or flipped alignments. However, by introducing slight asymmetric perturbations, we aimed to establish a global optimum with the lowest GWD among the local minima. In this case, the global minimum would involve diagonal correspondences (i.e., the mapping that aligns the same indices). We applied GWOT between these two circles to investigate which optimization process could identify the solution with the minimal GWD. We compared pairs of initialization strategies and epsilon samplers: “Random + TPE”, “Random + Grid Search”, and “Uniform + Grid Search”.

**Fig 6.**
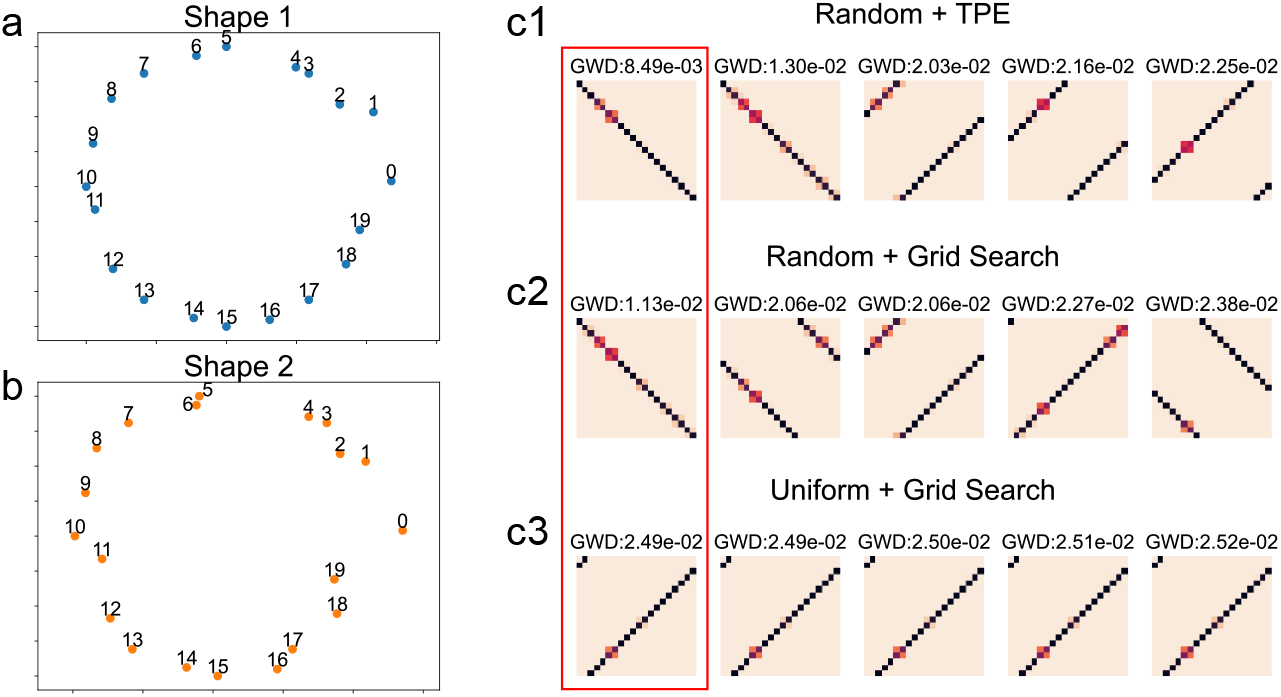
the impact of the initialization strategies. (a, b) Synthetic data was generated by first creating a circle with asymmetric perturbations (a), and then adding noise to create a second circle (b). Each point was assigned an index. (c1, c2, c3) Optimal transportation plans between shape 1 and shape 2 obtained through the three different strategies. The GWD values for each transportation plan are displayed above. For each strategy, the five transportation plans with the lowest GWD are shown. The matrix on the far left in the red square corresponds to the transportation plan with the lowest GWD.

As we expected, using “Random” initialization, GWOT identified the diagonal transportation plan, which corresponds to the global optimum, achieving a GWD of 0.0085 with TPE and 0.011 with grid search (Fig. 6c1, c2). In contrast, with a “Uniform” initialization, the optimal solution was not the diagonal matrix (the global optimum solution) and its GWD value is 0.025, which is much higher than the global optimum, 0.0085 (Fig. 6c1, c3). These findings indicate that the random initialization successfully identified the global optimum by searching for broader ranges of transportation plans while the “Uniform” initialization converged on a non-optimal solution. This tendency is also reflected in the distribution of the local minima. In the “Uniform” setting, the solutions were nearly identical (Fig. 6c3), while both “Random + Grid Search” and “Random + TPE” converged on a variety of transportation plans (Fig. 6c1, c2). This suggests that random initialization not only aids in finding the global optimum but also facilitates the exploration of the distribution of local minima, which may better reflect the underlying structure of the data.

Below, we performed all the following analyses using several common settings. For hyperparameter tuning, we used the Tree-Structured Parzen Estimator (tpe| option) to sample *ε* values and used random initialization of transportation plans (random| option) (see the Supporting Information for details on these options). The matching rate of the optimal transportation plan found by GWOT is evaluated based on the assumption that the correct assignment matrix is the diagonal matrix; in other words, that the indices between two dissimilarity matrices are aligned in the same order, and thus, the same indices correspond to each other.

### Synthetic data illustrating the differences between supervised and unsupervised comparison

Here, we show that unsupervised comparison can reveal structural similarities or differences between distance structures with greater detail than supervised comparison (conventional RSA). As mentioned above, in neuroscience, the similarity of distance structures is conventionally assessed using Representational Similarity Analysis (RSA) [3, 26, 27]. In the conventional RSA framework, the similarity of distance structures is typically evaluated using Pearson or Spearman correlation between distance matrices. A high correlation coefficient obtained through RSA indicates that the two representational similarity structures are alike. However, as shown in Fig. 3, even when there are high correlations between the representational structures, different scenarios can arise in terms of unsupervised alignment. In the following, we present three examples that illustrate the qualitative differences between our unsupervised and supervised approaches to comparing the structures.

First, we examine the case where the supervised and unsupervised comparison methods lead to qualitatively similar conclusions (Fig. 7(a)). Specifically, we constructed an example where we expect items in two structures to correspond one-to-one (Figs. 3(a), (b)). We first created two sets of circles and added correlated noise to both. Additionally, we introduced white noise to only the second set of circles. We then calculated the dissimilarity matrices for each set (Fig. 7(a1)). In the supervised comparison, the correlation coefficient between the two dissimilarity matrices in Fig. 7(a1) is very high (0.9). Fig. 7(a2) shows the optimal transportation plan, Γ, found by applying the GWOT method to the two dissimilarity matrices. In this case, the diagonal elements of Γ have high values, indicating that neural responses to the same stimuli are matched between the different brains, as depicted in Fig. 3(b). The unsupervised comparison method similarly confirms that the two dissimilarity structures are closely aligned at the level of individual stimuli. This conclusion is consistent with that obtained using the supervised comparison method.

**Fig 7.**
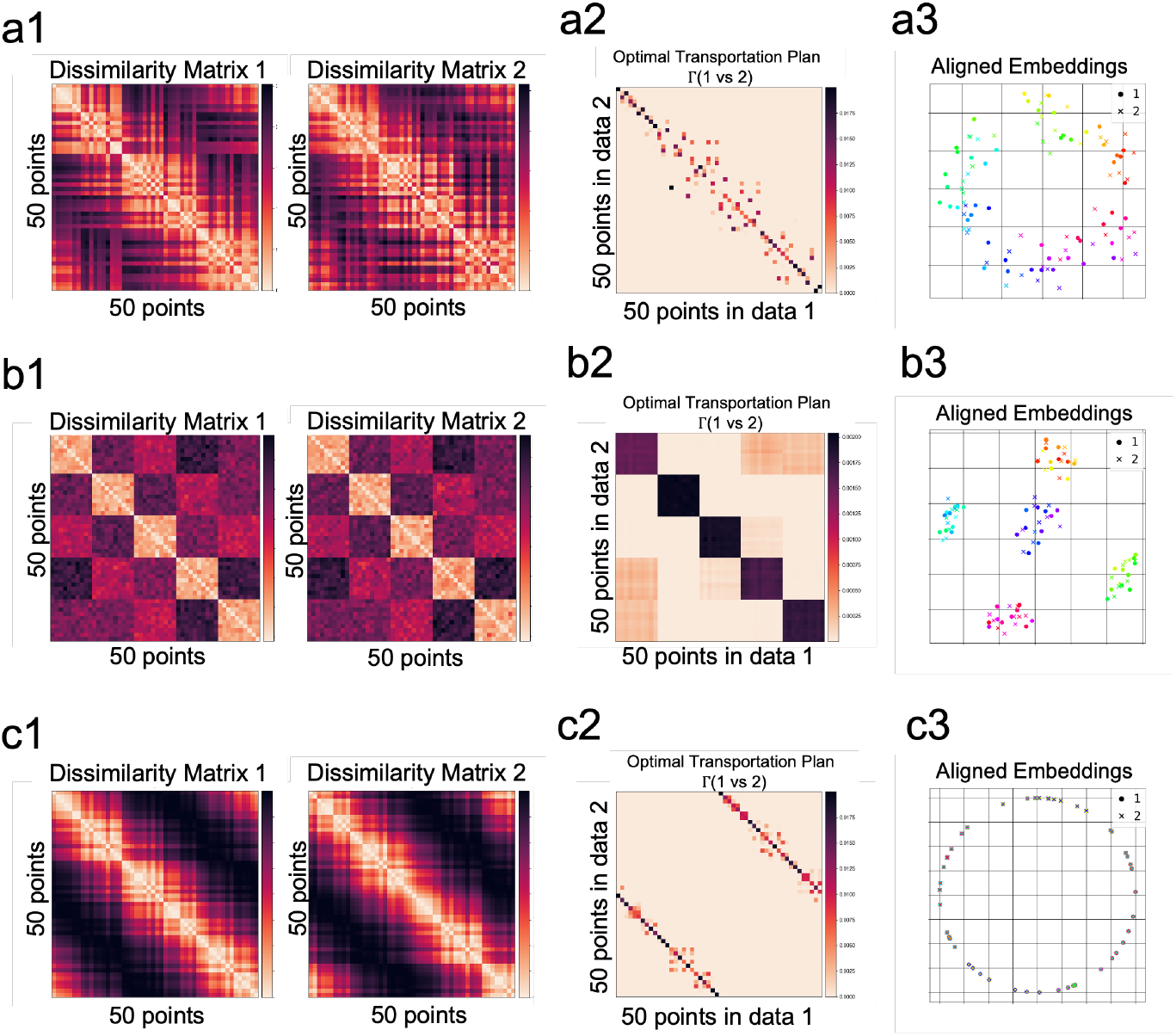
Unsupervised alignment in synthetic data, revealing structural similarities and differences with greater detail than supervised alignment. In all three cases, the Pearson correlation based on the conventional RSA framework (we call this “supervised comparison”) between the two distance matrices is constant at 0.90. In contrast, unsupervised alignment yields qualitatively different results, distinguishing the structural differences between the three cases. (a1, b1, c1) Two distance matrices of 50 points in the two groups. (a2, b2, c2) Optimized transportation plan Γ^*^ between the dissimilarity matrices 1 and 2. (a3, b3, c3) Embeddings of dissimilarity matrix 1 (*X*) and the aligned embeddings of dissimilarity matrix 2 (*QY*) plotted in the embedded space of *X*. (a) Case in which there are one-to-one level correspondences between the embeddings of *X* and *Y*, and that unsupervised alignment correctly finds the similar correspondences. (b) Case in which there are categorical (many-to-many) level correspondences between *X* and *Y*. (c) Case in which the assumed “correct” correspondences differ from the correspondences empirically found by unsupervised comparison.

However, the conclusions drawn from supervised and unsupervised comparisons can sometimes differ qualitatively (Figs. 7(b), (c)). For instance, the optimal correspondences between two structures may occur only at a broad categorical level, as illustrated in Fig. 3(c). In this example, we designed a scenario where we expect the items in the two structures to correspond at the coarse category level. We created sets of points to represent the existence of broad categories within the stimulus set (e.g., dogs, cats, cars, etc.), which are reflected as block structures in the dissimilarity matrix (Fig. 7(b1)). Although the correlation coefficient remains high (0.9), as in Fig. 7(a), the unsupervised comparison reveals that the alignment occurs only at the level of coarse categories, with no matching at the level of individual stimuli (Fig. 7(b2)).

Alternatively, items in two structures may exhibit shifted correspondences, as illustrated in Fig. 3(d). In this example, we generated two circles with a small augmentation and then rotated the labels of each point in the second circle, effectively shifting the identity of the stimuli (Fig. 7(c1)). In this case, we expect the two structures to have shifted correspondences while maintaining similar appearances. As a result, although the correlation coefficient remains high (0.9), as in Fig. 7(a), the unsupervised comparison reveals these shifted correspondences, indicating that neural responses to certain stimuli in one brain may correspond to those of “different” stimuli in the other brain (Fig. 7(c2)). This contrasts with the scenario in Fig. 7(a), where the neural responses to the same stimuli align. Figs. 7(a3), (b3), and (c3) show the aligned embeddings, highlighting the underlying structural differences across the three cases.

Taken together, unsupervised comparison reveals a more detailed structural correspondence between two sets of embeddings, whether it is a one-to-one fine “correct” mapping (Fig. 7(a)) or a many-to-many more coarse mapping (Fig. 7(b)), or the mapping is different from the assumed correspondences (Fig. 7(c)). This is clearly and qualitatively different from the supervised comparison method because it gives exactly the same evaluation in the three cases, and thus cannot distinguish the results of Figs. 7(b) and (c) from the result of Fig. 7(a).

### Behavioral data: Human psychological embeddings of natural objects

To demonstrate the alignment of behavioral data, we used the THINGS dataset, an open dataset containing human similarity judgments for 1,854 naturalistic objects [17–19]. As an example, we examined whether the similarity structures of 1,854 objects in the male and female participant groups can be aligned with GWOT. We first estimated the psychological embeddings of 1,854 objects for the male and female participant groups (see Supporting Information for details). We then computed the dissimilarity matrix of the 1,854 natural objects for each participant group, in which the dissimilarity between objects is quantified by the Euclidean distance between the embeddings of the objects (Fig. 8(a)). As can be seen in Fig. 8(a), the two dissimilarity matrices are closely similar, with a correlation coefficient of 0.96. However, as we showed in the toy example (Fig. 7(b)), a high correlation does not necessarily mean that the dissimilarity structures between the two groups match each other at the individual stimulus level. In fact, in the THINGS dataset, there are also coarse categories in 1,854 objects, and thus it is possible that the two structures only match at the coarse categorical level, as shown in 7(b).

**Fig 8.**
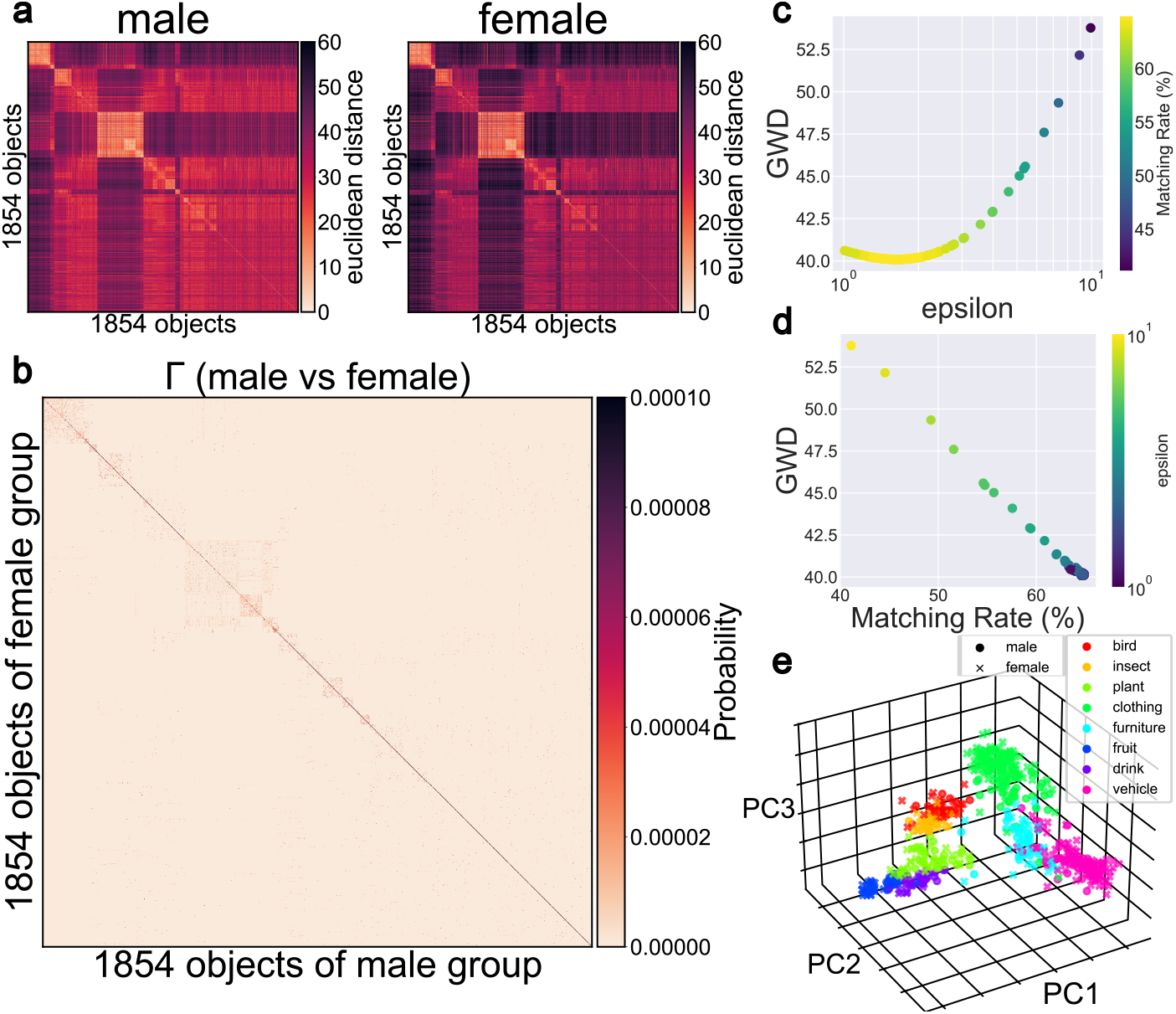
Unsupervised alignment between the psychological embeddings of naturalistic objects in two different participant groups. (a) Dissimilarity matrices of 1,854 naturalistic objects in the male and female participant groups. (b) Optimized transportation plan Γ^*^ between the distance matrices of the male and female groups. (c) Relationship between GWD and hyperparameter *ε*. Color represents matching rate. (d) Relationship between GWD and matching rate. Color represents hyperparameter *ε*. (e) The embeddings of the male group and the aligned embeddings of the female group plotted in the embedded space of the male group. Only objects with certain coarse category labels (“bird”, “insect”, “plant”, etc.) are shown here. Each point represents the embedding of an object. The colors represent the coarse categories of the objects.

Next, we performed GWOT between the dissimilarity matrices of the male and female participant groups. In Fig. 8(b), we show the optimized transportation plan with the minimum GWD between the male and female participant groups. As shown in Fig. 8(b), most of the diagonal elements in Γ^*^ have high values, indicating that most of the objects in one group correspond to the same objects in the other groups with high probability. This result means that the two similarity structures match at the level of individual objects.

To show the optimization results in detail, we reveal the relationship between the hyperparameter *ε*, GWD, and the matching rate of the unsupervised alignment over 100 iterations in Figs. 8(c) and (d). Fig. 8(c) shows that good local minima with a low GWD as well as high matching rate are found over a wide range of epsilon between 1 and 10. Fig. 8(d) shows a downward trend to the right, i.e., lower GWD values tend to result in higher accuracy. This downward trend to the right is necessary for successful unsupervised alignment. The optimal solution with the minimum GWD has the matching rate of 64.7%.

Finally, using the optimal transportation plan, we aligned the psychological embeddings of natural objects across the two groups of participants. For visualization, we projected the original embeddings in 90 dimensions into three dimensions using Principal Component Analysis (PCA) (Fig. 8(e)). Also, since the display of all 1,854 objects would result in crowding, we show only those objects belonging to the 8 categories indicated by different colors in Fig. 8(e). We can see that the projected neural responses to the same categories are positioned close to each other in the projected space. The top-1 categorical level matching rate is 85.5%.

### Neural data: Neuropixels visual coding in mice

To demonstrate the alignment of neural activity data, we used the Neuropixels Visual Coding dataset from Allen Brain Observatory (see Supporting Information for details). First, we created two “pseudo-mice” by dividing 30 mice into two non-overlapping groups and aggregating the neural responses of 15 mice in each group. Second, for each “pseudo-mouse”, we computed the dissimilarity matrix of the neural responses of higher dorsal visual cortex, anterolateral visual cortex (VISal), and anteromedial visual cortex (VISam), to 90 segmented short movie stimuli (Fig. 9(a)). The dissimilarity is quantified as the cosine distance between the neural responses of the pseudo-mice. As can be seen, these two dissimilarity matrices are quite similar (correlation coefficient *ρ* between them is 0.92).

**Fig 9.**
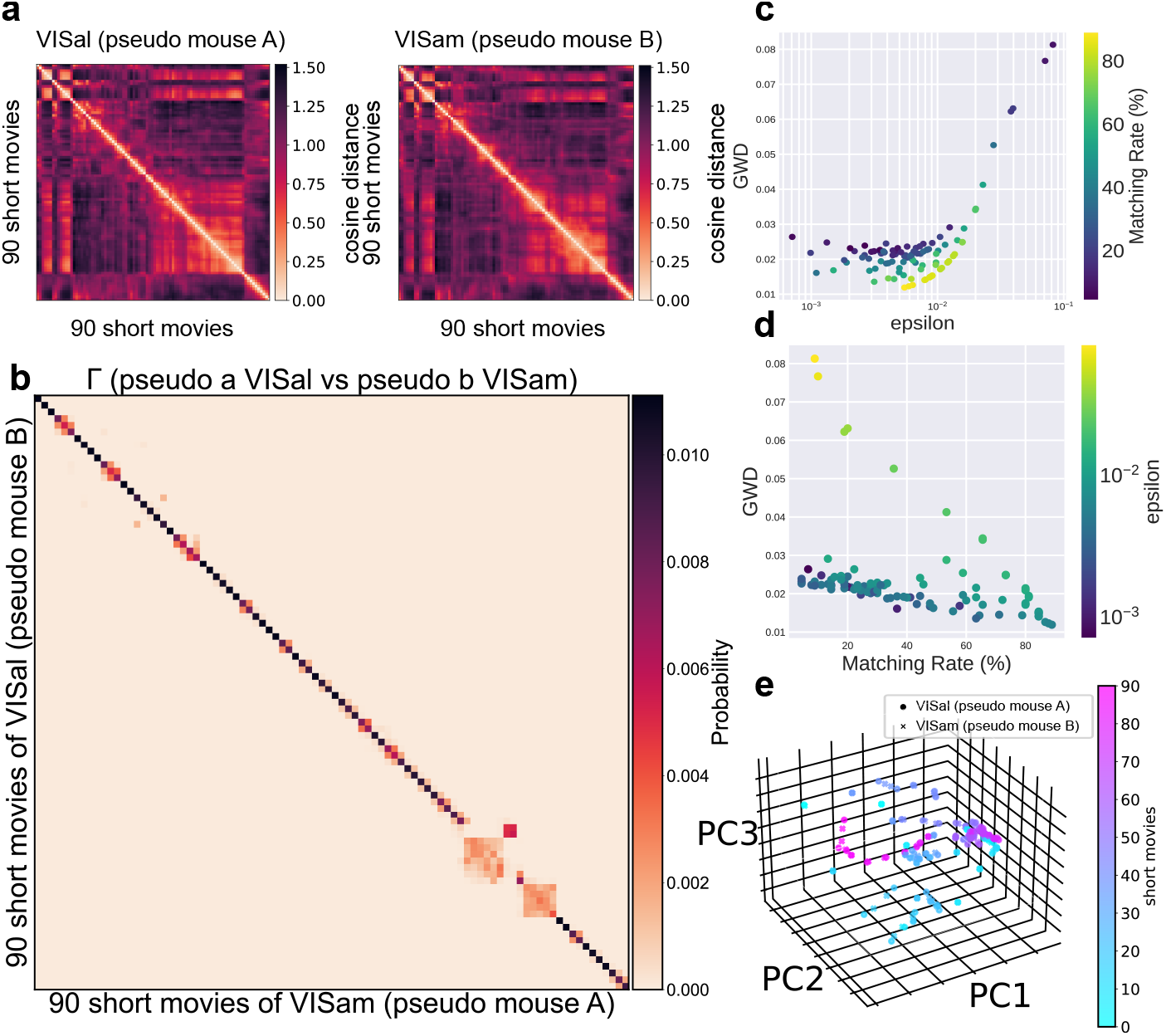
Unsupervised alignment between the neural representations of anterolateral visual cortex (VISal) and those of anteromedial visual cortex (VISam) in different two pseudo-mice. (a) Dissimilarity matrices in the two pseudo-mice for 90 segmented short movies, made from a 30-second continuous natural movie stimulus. (b) Optimized transportation plan Γ^*^ between the dissimilarity matrices of the pseudo-mice. (c) Relationship between GWD and hyperparameter *ε*. Color represents matching rate. (d) Relationship between GWD and matching rate. Color represents hyperparameter *ε*. (e) The embeddings of pseudo-mouse A and the aligned embeddings of pseudo-mouse B plotted in the pseudo-mouse A’s embedded space. Colors represent the number of the short movie stimuli.

Next, we performed the optimization of GWD between the dissimilarity matrices of the different areas in the two pseudo-mice. To optimize the hyperparameter *ε*, we sampled 100 different values of *ε* within a range of 10 ^−4^ to 10 ^−1^. For each sampled *ε*, we randomly initialized the transportation plans. We show the optimized transportation plan with the minimum GWD in Fig. 9(b). As can be seen, almost all of the diagonal elements have the highest values (top-1 matching rate is 92.2%). Even in the case of mismatch of movie segments, the mismatch occurs between the movie segments that are close in time and thus similar to each other. This demonstrates a strong structural correspondence between the neural responses in the dorsal visual cortex.

To see the details of the optimization results, we show the relationship between the hyperparameter *ε*, GWD, and the matching rate of the unsupervised alignment over 100 iterations in Figs. 9(c) and (d). Fig. 9(c) shows that GWD is optimized when *ε* is around 10 ^−2^. Fig. 9(d) shows a downward trend to the right, i.e., lower GWD values tend to result in higher accuracy. This downward trend to the right is necessary for successful alignment.

Finally, using the optimal transportation plan with the minimum GWD, we aligned the neural responses of the two pseudo-mice. We performed this alignment using the responses of all neurons in each pseudo-mouse. For visualization, we projected the 738 dimensional neural responses into three dimensions using Principal Component Analysis (PCA) (Fig. 9(e)). We can see that the projected neural responses to the same movie stimuli are positioned close to each other and also that the responses to movie stimuli close in time (indicated by colors) are close to each other in the projected space.

### Model: Vision Deep Neural Networks

To demonstrate the alignment of neural network models, we used two vision deep neural network models, ResNet50 and VGG19 (see Supporting Information for details). First, we computed the dissimilarity matrices of the 1,000 natural images belonging to 20 classes for ResNet50 and VGG19 as the cosine distance between the last fully connected layer of the two DNNs. (Fig. 10(a)). The correlation coefficient between the two matrices are fairly high, with a correlation coefficient of 0.91. However, since there are 20 image classes, this high degree of correlation can be induced simply by categorical level correspondences, as illustrated in the toy example in Fig. 7(b) (note that the rows and columns of the matrices are sorted by class labels, i.e., 50 images from index *i* to index *i* + 49 belong to the same class where *i* ≡ 0 mod 50). Thus, it is important to assess whether these embeddings match at the individual image level or merely at the categorical level by using the unsupervised alignment.

**Fig 10.**
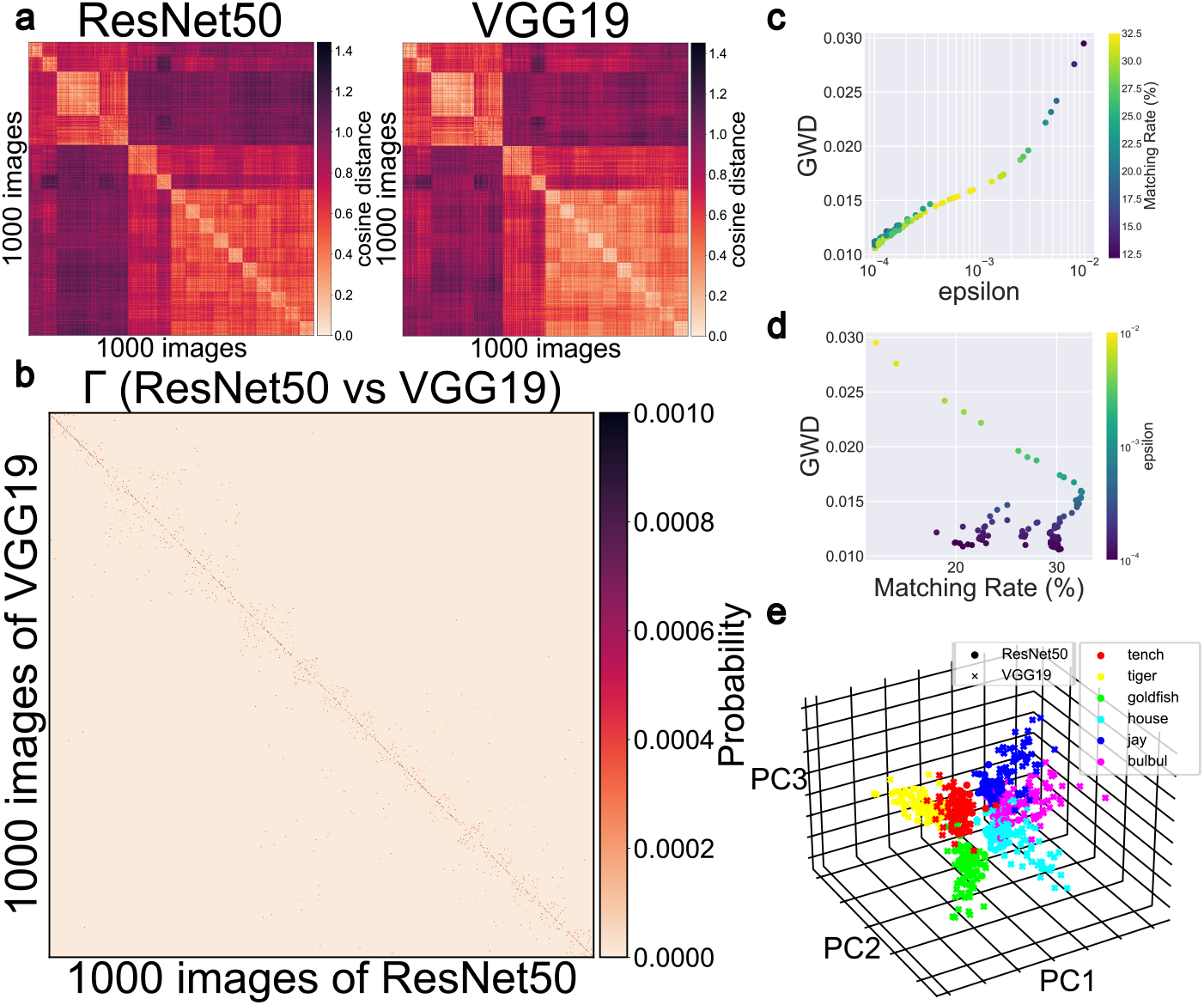
Unsupervised alignment between embeddings of the fully-connected layers of the vision DNNs, ResNet50 and VGG19. (a) Dissimilarity matrices of 1,000 natural objects in the two different vision DNNs, ResNet50 and VGG19. (b) Optimized transportation plan Γ between the dissimilarity matrices of ResNet50 and VGG19. (c) Relationship between GWD and hyperparameter *ε*. Color represents matching rate. (d) Relationship between GWD and matching rate. Color represents hyperparameter *ε*. (e) Embeddings of ResNet50 and the aligned embeddings of VGG19 plotted in ResNet50’s embedded space. All objects with a class label (“goldfish”, “house”, “tiger”, etc.) are shown. Each dot represents the embedding of each object. Colors represent the class of the images.

Next, we performed the optimization of GWD between the dissimilarity matrices of the two DNNs. As shown in Fig. 10(b), many diagonal elements in Γ^*^ have high values, indicating that many embeddings in one DNN correspond to the embeddings of the same images in the other DNN with high probability. This result means that the two similarity structures of the two DNNs match at the level of individual images, even though the details of the network architectures are largely different. Even for mismatches, we can see high values close to the diagonal elements, which means that the embeddings in one DNN match those belonging to the same image class in the other DNN because the image indexes in Fig. 10(a) are sorted by class labels.

To see the details of the optimization results, we show the relationship between the hyperparameter *ε*, GWD, and the matching rate of the unsupervised alignment over 100 iterations in Figs. 10(c) and (d). Fig. 10c shows the local minima with lower GWD values in the lower epsilon ranges below 10 ^−3^. Fig. 10(d) shows a downward trend to the right, i.e., lower GWD values tend to result in higher matching rate, as is also the case in Fig. 8 and Fig. 9. The optimal solution with the minimum GWD has a top-1 matching rate of 30.2%.

Finally, using the optimal transportation plan, we aligned the embeddings of the two vision DNNs. For visualization, we projected the original embeddings of 1,000 model neurons into three dimensions using Principal Component Analysis (PCA) (Fig. 10(e)). Also, since the display of all 1,000 objects would lead to crowding, we show only the objects belonging to 6 image classes, indicated by different colors in Fig. 10(e). We can see that the projected neural responses to the same categories are positioned close together in the projected space. The top-1 categorical level matching rate is 78.1%.

## Summary and future directions

In this paper, we present a comprehensive guide to unsupervised alignment based on GWOT to facilitate many possible use cases, especially for neuroscience, but also for other research areas. We also provide an effective and user-friendly toolbox for hyperparameter tuning of entropic GWOT. The source code for our toolbox is available at https://oizumi-lab.github.io/GWTune/. All examples shown in this paper can be tested in the tutorial notebooks to help new users get started. In addition, another example using the similarity judgment data of 93 colors previously reported in [5] is also available. The processed data used to generate the results presented in this study are also available in the “data” folder of the same GitHub repository.

An important variant of the GWOT, not covered in this paper and our toolbox, is called the partial or unbalanced GWOT [31, 32]. The normal or balanced GWOT is known to be vulnerable to the presence of outliers. The unbalanced GWOT is a more flexible method than the balanced GWOT because it can ignore outliers and try to find a partial mapping between similarity structures. Such methods would be necessary when the similarity structures to be compared are not entirely the same as a whole structure, but are only partially equivalent. We have not yet implemented it in GWTune due to the lack of a clear methodology for hyperparameter tuning in this context. The presence of an additional hyperparameter, *ρ*, which controls the overall mass, complicates the selection of optimal hyperparameters based solely on the minimum GWD criterion. We are currently investigating effective hyperparameter tuning strategies for unbalanced GWOT, and plan to incorporate this functionality into GWTune in a future update, leveraging the advanced hyperparameter optimization capabilities of Optuna.

Finally, we discuss several important cases for the application of GWOT that are not covered in this paper. The first is the case in which the number of points (embeddings) is different. In this study, we only consider the case where the number of points is the same and the set of external stimuli or inputs is the same. For example, in the case of the Allen Neuropixels data, the same movie stimuli are used between the pseudo-mice.

However, GWOT is applicable to cases in which the number of stimuli is different, or the stimuli themselves are different. For example, we can consider the alignment between neural responses to a set of *N* stimuli in one brain and neural responses to a different set of *M* stimuli in the other brain. GWOT can be readily used in such cases and will be useful in finding the correspondences among neural responses to different stimuli.

The second case involves the alignment between data from different modalities. For example, we can consider the alignment between neural activity and behavior or a model. This application is important for assessing how well a given neural network model captures neural responses in the brain or how well it explains behavior in humans or other animals [33–38]. Moreover, we can also consider the alignment between behavior and neural activity, which is important for identifying neural correlates that explain a given behavior in humans or other animals [19, 39–44]. Although this paper focuses primarily on aligning representations within a single modality, GWTune is also capable of handling different modalities. As an example, our group applied this toolbox to evaluate whether human behavioral similarity data can be unsupervised aligned with similarity data from neural network models [12, 13]. We hope that this toolbox will assist researchers in applying GWOT to various types of data, including cases such as these, and provide novel insights that cannot be obtained by conventional supervised alignment methods.

## Acknowledgments

We thank Genji Kawakita for early code contributions. We also thank Ariel Zeleznikow-Johnston and Naotsugu Tsuchiya for providing the data on color similarity judgments for the toolbox tutorial. We thank the authors of the THINGS data for providing us with early access to the THINGS data. This work was supported by JST Moonshot R&D Grant Number JPMJMS2012, and JSPS KAKENHI Grant Numbers 20H05712 and 23H04834.

## Supporting Information

This Supporting Information includes the following.

- Alignment of point clouds
  – Supervised alignment
  – Unsupervised alignment
  – Unsupervised alignment by barycentric projection
- Evaluation of the optimization results of GWOT
  – Matching rate of optimal transportation plans
  – Inspection of other local minima
  – Visualization of aligned embeddings
- Details of code and implementation
  – Software Requirement
  – Parameters for the optimazation of entropic GWOT
    – Hyperparameter tuning of *ε*
    – Initialization of transportation plans Γ
  Comparison between previous existing software and our toolbox
- Data and pre-processing
  – THINGS data
  – Neuropixels visual coding from the Allen Brain Observatory
  – Vision deep neural network models
- Comparison of different initialization strategies with three real-world datasets

### Alignment of point clouds

Here we consider the problem of aligning two sets of point clouds *X* and *Y* (Figs. 2b and c). These points represent, for example, neural responses, behavioral responses, or activity in a neural network model to certain stimuli. We commonly call these points “embeddings”. *X* and *Y* are *d* × *n* matrices, where *n* is the number of embeddings and *d* is the dimension of embedding vectors.

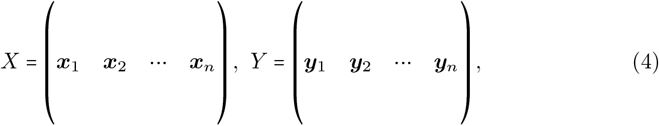

where ***x***_*i*_ and ***y***_*i*_ are column vectors representing the *i*th embeddings of *X* and *Y*, respectively.

The general problem setting in this study is to find the optimal alignment between *X* and *Y* without assuming any correspondence between the columns (the embeddings) of *X* and *Y*. For example, we may sometimes know that ***x***_*i*_ and ***y***_*j*_ are the neural responses to the same external stimulus, suggesting that the *i*-th column of *X* corresponds to the *j*-th column of *Y*. However, in the general unsupervised alignment setting, we do not use this information.

As a general problem, we consider solving the following problem:

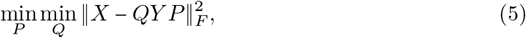

where ∥ ⋅ ∥_*F*_ is the Frobenius norm 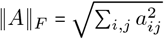, *P* is the *n* × *n* assignment matrix that establishes correspondence between the column vectors of *X* and those of *Y* (i.e., ***x***_*j*_ ↤ ∑_*i*_ *P*_*ij*_ ***y***_*i*_), and *Q* is the *d × d* orthogonal matrix that rotates *Y* to fit into *X*. If we restrict *P* such that each column has exactly one element equal to 1 and the others equal to 0, the problem becomes finding a one-to-one correspondence between the columns (embeddings) of *X* and *Y*, or equivalently, finding the optimal permutation of the column indices of *X*. However, in this study, we investigate a more general scenario where the elements of the matrix *P* can take any real value between 0 and 1. These values represent the degree of correspondence between the *i*-th column of the matrix *X* and the *j*-th column of the matrix *Y*. This more flexible approach allows us to model the correspondences between the columns (embeddings) of *X* and *Y* in a more comprehensive way.

There are two broad categories of methods for this general problem, supervised alignment and unsupervised alignment. These are explained in detail in the following sections.

#### Supervised alignment

Supervised alignment is a method in which the assignment matrix *P* is given. In the case of the optimization problem in Eq. 5, it becomes the well-known Procrustes problem [25], which has a closed-form solution. For example, if we simply assume that the column indices of *X* match those of *Y*, and therefore *P* is the identity matrix, the optimization problem is given by

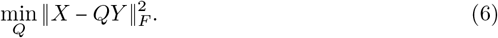

Given the singular valu^*^ e decomposition *U* Σ*V* ^⊺^ of *XY* ^⊺^, the solution to the Procrustes problem is given by *Q*^*^ = *UV*^⊺^.

#### Unsupervised alignment

Unsupervised alignment is the method where the assignment matrix *P* is not given. In this case, we need to jointly optimize *P* and *Q* in Eq. 5, which is a non-convex optimization problem without a closed-form solution.

One way to address this problem is to use the optimal transportation plan found in Gromov-Wasserstein Optimal Transport (GWOT) as the optimal assignment matrix *P*. Then, based on the assignment matrix, i.e., the optimal transportation plan, the Procrustes solution *Q*^*^ is computed. This approach has been used in some previous studies [8, 9]. In this paper, we also used this approach to visualize the unsupervised alignment based on GWOT.

Denoting the optimal transportation plan (the assignment matrix) by Γ^*^, the problem to solve becomes

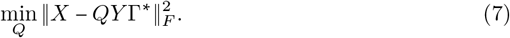

The solution can be found by the singular value decomposition of *X* (*Y* Γ^*^)^⊺^.

#### Unsupervised alignment by barycentric projection

Another method of unsupervised alignment based on GWOT used in some previous studies [10, 16] is a barycentric projection. While the method based on the Procrustes approach (Eq. 7) can only be applied if the dimensionality of the embeddings in *X* and *Y* is the same, or the dimensionality has to be adjusted to be the same before application, the barycentric projection method can always be used even if the dimensionality of the embeddings *X* and *Y* is different.

With the optimized transportation plan Γ found by GWOT, the barycentric projection is defined as

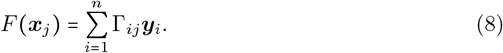

This mapping function *F* can be interpreted as projecting the embedding ***x***_*j*_ into the space of the embeddings *Y*. By applying this projection, all the mapped embeddings of *X*, (*F (****x***_1_), *F (****x***_2_), ⋯, *F (****x***_*n*_), and the embeddings of *Y*, (***y***_1_, ***y***_2_, ⋯, ***y***_*n*_) can be compared within the same space.

Although in this paper, we use the Procrustes approach because the dimensionality of the embeddings to be compared is the same for all the examples, one can consider using this barycentric projection approach for some applications.

### Evaluation of the optimization results of GWOT

#### Matching rate of optimal transportation plans

If there is a ground truth mapping or an assumed correspondence between two sets of embeddings, it is recommended to compute the correct matching rate between the empirically found correspondence and the known correspondence. One of the ways to compute the correct matching rate is based on optimal transportation plans. To define it, we define the “correct” assignment matrix *C* as *C*_*ij*_ = 1 if *i* and *j* are matched and *C*_*ij*_ = 0 if otherwise. By using the correct assignment matrix *C*, the correct pairs matching rate based on the optimal transportation plan is defined as follows. When *i* and *j* are a correctly matched pair, the following function checks if the element of the transportation matrix, Γ_*ij*_, is the maximum among the other elements Γ_*ij*_′ in the same row,

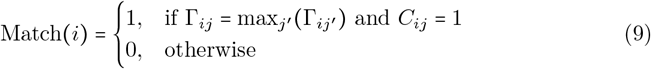

The matching rate is then the percentage of index *i* that matches with the correct pair *j*, which can be calculated as:

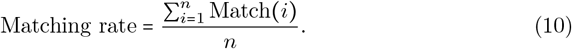

The matching rate defined above is the top-1 matching rate. More generally, the top-*k* matching rate is similarly computed by checking whether Γ_*ij*_ is within the top-*k* highest values among the other elements Γ_*ij*_′ when *C*_*ij*_ *=* 1.

Although the matching rate defined above is the evaluation of the matching at the fine-item level, in some datasets where the stimuli are classified into several coarse categories, we may want to evaluate the matching at the coarse-categorical level. To evaluate the extent to which items were matched with the items belonging to the same category, but not necessarily the exact correct pairs, we can use the measure “categorical level matching rate”, defined as follows. When *i* belongs to a category *G*, the following function checks whether the element *j* that receives the maximum amount of transportation from the element *i* is in the same category *G* as *i*:

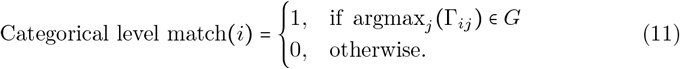

The categorical level matching rate is then defined as the percentage of indices that match any of the indices within the same category:

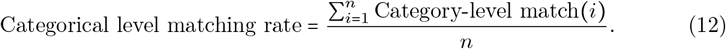

#### Inspection of other local minima

It is crucial and highly recommended to not only focus on the best optimal transportation plan with the minimum GWD value, but also to examine the other local minima with higher GWD values. First, it is always helpful to plot the relationship between GWD and *ε* to investigate how GWD values depend on *ε* and how close or far the best solution with the minimum GWD value is from other local minima.

Second, when evaluating the matching rate, it is also informative to plot the relationship between the GWD values and the matching rate to understand whether a high matching rate is likely to be robustly achievable. For example, if it is observed that the optimal transportation plans with lower GWD values tend to result in higher matching rates, and the GWD values of the optimal transportation plans with high matching rates are far enough away from those with low matching rates, then it would imply that a high matching rate is likely to be achievable.

#### Visualization of aligned embeddings

It is helpful to visually inspect the quality of the unsupervised alignment based on GWOT by plotting the aligned embeddings in 2D or 3D space. Here, we consider the alignment of the embeddings of *Y* with the embeddings of *X*. By multiplying *Y* by the rotation matrix *Q*^*^ obtained from the Procrustes analysis in Eq. 7, we obtain the aligned embeddings *Q*^*^ *Y*. Then, by plotting *Q*^*^ *Y* and *X* (the pivot embeddings) in the same space, we can compare these two embeddings and evaluate the quality of the unsupervised alignment. For visualization, Principal Components Analysis (PCA) is for example used to project high-dimensional embeddings into 2D or 3D space, but other dimensionality reduction methods are also applicable. If there is an assumed correspondence between the embeddings, one can visually check whether the aligned embeddings are positioned close to the corresponding embeddings.

### Details of code and implementation

#### Software Requirement

Our toolbox can be run and tested in Python version ≥3.9. Important packages used in our toolbox are POT and Optuna. The compatible versions are POT version 0.9.4 and Optuna version ≥ 3.2. The compatible versions of other packages are described in the ReadMe in the GitHub repository (https://oizumi-lab.github.io/GWTune/).

#### Parameters for the optimazation of entropic GWOT

##### Hyperparameter tuning of *ε*

To tune the hyperparameter *ε* values, the range of *ε* values, how many trials *ε* values are sampled, and the sampling method need to be set. Setting an appropriate range for *ε* is crucial for the efficient performance of GWOT, as values that are too high or too low may not produce optimal results. While a wide range of *ε* can still yield good values if the algorithm runs over many trials, it is generally advisable to narrow down the range after conducting a small number of initial test runs, before performing a more extensive search.

As for sampling *ε*, the following three sampling methods, originally implemented by Optuna, are available in the toolbox. First, `tpe`, an efficient sampler based on Bayesian sampling, is generally recommended for most cases, and the availability of such efficient sampling methods is indeed the reason why Optuna is used in the toolbox. tpe is the default sampler in Optuna, which has been shown to perform well in many settings (see https://github.com/optuna/optuna/issues/2964 for details of the benchmark experiment). The other two samplers are simple grid search and random search samplers. In most cases, tpe will outperform these alternatives because it leverages past trial results to identify ranges of *ε* values that are more likely to achieve lower GWD, whereas grid and random search sample *ε* values independently of previous trials. See [15] for the details of each sampler.

`tpe` (recommended): A sampler that uses Tree-structured Parzen Estimator (TPE) based on Bayesian optimization. Given the past history of the values of the objective function *y* (GWD in our case) under hyperparameter values *x*, the estimator attempts to maximize or minimize the expected improvement of *y* by modeling the conditional probability, *p (x* ∣*y)*, based on kernel density estimation. See [45, 46] for mathematical details.

`grid:` A grid search sampler that exhaustively samples *ε* values on a user-specified grid in a one-dimensional *ε* space.

`random:` A sampler that randomly and independently samples *ε* values from the user-defined range of *ε* values.

##### Initialization of transportation plans Γ

There are several initialization methods for transportation plans. All of them ensure that the initial transportation plans Γ satisfy the mass conservation constraint, i.e., the marginals of the transportation plans are equal to ***p*** and ***q***.

A default method (`random`) is to initialize the transportation plans randomly and independently on each trial, while the other methods are fixed initialization over all trials. In general, trying many different initial values is more likely to find better local minima than trying only one fixed initial value. Thus, we recommend the random option over fixed initialization for most use cases.

The `diag` option or the `user_define` option should be used with care in unsupervised alignment to avoid “cheating”. In the strictest sense, unsupervised alignment is a problem setting where alignment is searched without relying on known or presumed correspondences. In this strict problem setting, use of the `diag` option or `user_define` option only is considered cheating because these options inherently tend to find local minima with a high matching rate in unsupervised alignment. When using these options, the following must be done. First, the `random` option should also be used, and the local minima found by the random option should be compared fairly with those found by some given correspondences. If the `random` option finds local minima with lower GWD but a lower matching rate, one must consider these local minima as the optimal solutions found by the GWOT algorithm instead of those found by the `user_define` or `diag` options. Second, the fact that some correspondences are given should be explicitly stated to distinguish results with some known information from those without any given information.

Note, however, that the use of the `diag` or `user_define` option is justified or even preferred in problem settings other than the strict unsupervised alignment. For example, in the semi-supervised alignment problem setting, where some of the correspondences are known but the others are not, providing the known correspondences as an initialization will be a natural choice.

`random` (recommended): This method initializes each element of the transportation matrix by sampling from a uniform distribution [0, 1], and then normalizing it to satisfy the mass conservation constraint. The number of initialization on the same trial with the same *ε* value can also be specified. Note, however, that even if the number of initialization on the same *ε* value is set to 1, if many different *ε* values are sampled, many different initialization will be performed on similar *ε* values when tpe is used. This is because tpe will extensively search for likely good *ε* values that will end up with similar *ε* values in later trials. Thus, increasing the number of initialization is similar to increasing the number of *ε* samples when using `tpe`.

`uniform`: This method initializes a transportation plan by taking the direct product of the marginals ***p*** and ***q***, i.e., ***p*** ⊗ ***q***. When both ***p*** and ***q*** are set to a uniform distribution, this method generates a uniform transportation plan consisting of uniform values, whereas if ***p*** or ***q*** is not a uniform distribution, the initial transportation plan is not a uniform matrix. However, for ease of understanding, we simply call this option `uniform`, assuming that ***p*** and ***q*** are set to a uniform distribution for most use cases. This method is the same as the default initialization implemented in POT (Python Optimal Transport) [30].

`diag` (use with caution): This method uses a diagonal matrix with constant values as initialization. This is intended only for the special case where there is a known one-to-one correspondence between the indices of the dissimilarity matrices and these indices are sorted in the same order. Other more general correspondence, such as one-to-many or many-to-many correspondence, should be provided by using the `user_define` option.

`user_define` (use with caution): This method initializes a transportation plan with a user-defined matrix. As in the case of “diag”, it is designed for cases where known or presumed correspondences can be provided.

#### Comparison between previous existing software and our toolbox

Our toolbox differs from existing software (e.g., POT [30], SCOT [10]) in two key aspects. First, in the optimization of GWOT, we offer a more flexible approach to setting the initial values for the transport plan. While existing software uses only a uniform distribution, we have added the option to use random initial values. Second, in tuning the hyperparameter related to entropy regularization, existing software usually relies on a fixed value or a grid search. In contrast, we have integrated an Optuna-based Tree-structured Parzen Estimator (TPE) sampler, in addition to the conventional methods (see Tab.1).

**Table 1.** Comparison between previous existing software and our toolbox.

|  | POT [30] | SCOT [10] | Ours |
| --- | --- | --- | --- |
| Initialization | Uniform | Uniform | Uniform / Random |
| Tuning of $\epsilon$ | Fixed | Grid Search | Grid Search / TPE |

### Data and Pre-processing

#### THINGS data

In the THINGS dataset, participants performed an odd-one-out task, where they were presented with three naturalistic objects from the THINGS dataset and asked to report which item in the triplet was the most dissimilar to the other two objects. This dataset includes approximately 4.70 million similarity judgments from about 12,000 participants collected through online crowdsourcing. To conduct the alignment between male and female participant groups, we randomly sampled an equal number of judgments from all male and female participants to create the male and female groups. Each participant group contains 1 million similarity judgments, a sample size shown to be sufficient for estimating meaningful and consistent representations of natural objects [18, 19]. After making the two participant groups, we next need to estimate the embeddings of the 1,854 natural objects from the odd-one-out judgment data. We followed the procedure of previous studies to estimate the embeddings [5, 18, 19].

First, the embeddings of the 1,854 objects were initialized with 90 randomly assigned dimensions ranging from 0 to 1. Second, the Euclidean distance between all pairs of the embedding vectors was computed and considered as the dissimilarity between the embeddings,

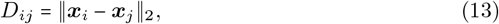

where ∥ ⋅ ∥_2_ is the L2 norm 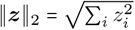. Conversely, the similarity between the embeddings was quantified as the negative Euclidean distance,

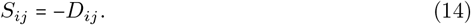

Third, using the similarity between the embeddings, we estimated the probability that a participant chooses image *k* as an odd object among the triplet *i, j, k*, which is equivalent to the probability of choosing image *i* and *j* as the most similar object pair among three possible pairs. Here, the probability was estimated by the softmax function of the similarity between the embeddings of the pair (*i, j)*,

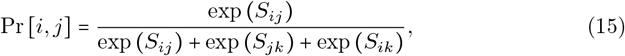

where *S*_*ij*_ is given by Eq. 14. Fourth, we updated the embeddings by minimizing the following loss function. For the *l*-th triplet in the dataset, let 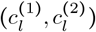 denote the index pair chosen by a participant as the most similar pair. Then, the loss function is given by

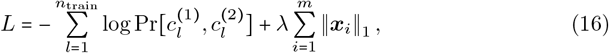

where *n*_train_ is the total number of triplets in the training dataset, *m* is the number of the natural objects, and ∥ ⋅ ∥_1_ denotes the L1 norm ∥***z***∥ _1_ = ∑*i* |*z*_*i*_|. The first term is the cross-entropy loss and the second term is the L1 norm regularization with the hyperparameter *λ*. The loss function was optimized by the Adam algorithm with an initial learning rate of 0.001, using a fixed number of 1,000 epochs. The hyperparameter *λ* was optimized by 5-fold cross-validation.

#### Neuropixels visual coding from the Allen Brain Observatory

The Neuropixels Visual Coding dataset consists of large-scale electrophysiological recordings in the mouse visual system including multiple cortical areas [20]. Among the recorded areas, we chose the anterolateral visual cortex (VISal) and the anteromedial visual cortex (VISam), which are parts of the higher dorsal visual cortex, as examples (see [14] for other areas). The number of neurons recorded in these areas is about 50, depending on the mouse. As for visual stimuli, we chose a natural movie stimulus (“natural movie one”), which is a 30-second scene from a black and white movie, as an example.

We considered the alignment of the aggregated neural responses across multiple mice, rather than the alignment of neural responses between individual mice. This is because we found that the alignment of individual mice was impossible for any of several possible reasons, such as the limited number of recorded neurons, fluctuation or noise in neural responses, and individual differences. Specifically, we aggregated the neural responses of 15 mice, corresponding to the responses of approximately 700 neurons in total, and considered them to be the neural responses of a “pseudo-mouse”. Similarly, we also made the aggregated neural responses from 15 different mice and considered it as another “pseudo-mouse”. We then considered the alignment between the two pseudo-mice created in this way. For visual stimuli, we used a natural movie stimulus (“natural movie one”). Although the movie stimulus is a 30-second continuous movie stimulus, we segmented it into 1/3-second short movie stimuli and treated them as 90 segmented movie stimuli.

After the standardization of the spike counts for each neuron, we then computed the trial average of the spike counts during each 1/3-second short movie stimulus and considered these as the neural responses of the pseudo-mice to be aligned.

#### Vision deep neural network models

For vision deep neural network models, we used the trained models of ResNet50 [22] and VGG19 [23] available in PyTorch as an example. We extracted the embeddings from the last fully connected layers in the two DNNs, whose dimensions are 1,000.

As input images, we used the validation set in the ImageNet dataset [47], which contains 50,000 natural images belonging to 1,000 classes. To simplify the computation, we used only 1,000 images belonging to 20 classes, where each class contains 50 images.

### Comparison of different initialization strategies with three real-world datasets

In addition to the theoretical consideration and simulations in artificial data in “Synthetic data illustrating the impact of the initialization strategies”, we also assessed whether the differences are also observed in real-world data. We systematically compared the change in the minimum GWD value over 100 trials for three strategies, `“Random + TPE|”, “Random + Grid Search|”`, and `“Uniform + Grid Search|”`, in three real-world datasets: THINGS, AllenBrain, and DNN. For each strategy, we sampled epsilon 100 times, depending on whether the sampling strategy was `“TPE|”` or `“Grid Search|”`. The range of epsilon values for each dataset is the same as those in the main results. Our results showed that both `“Random + TPE|” and “Random + Grid Search|”` strategies were consistently more effective at minimizing GWD than “Uniform + Grid Search|” across all datasets (Fig. S1). In particular, for the AllenBrain dataset, we observed that there is a significant difference between the initialization methods in terms of both the minimized GWD and the optimal transportation plan. With `“Random|”` initialization, GWOT converged to the diagonal transportation plan (Fig. S2b1, b2), achieving GWD = 0.012 and matching rate = 86.7% with TPE and GWD = 0.013 and matching rate = 76.7% with Grid Search (Fig. S1). In contrast, with “Uniform|” initialization, we observed in the optimal transportation plan that several mismatches occur, i.e., some movie chunks are mapped to other chunks in reverse time order (upward to the right) (Fig. S2b3), resulting in higher GWD (GWD = 0.015) and lower matching rate (matching rate = 63.3%) than the random initialization methods (Fig. S1). The differences between `“Random|”` and `“Uniform|”` are also evident in the optimization process (Fig. S1b), where `“Random + TPE|” and “Random + Grid Search|”` quickly minimize the GWD, while the GWD of `“Uniform + Grid Search|”` remains consistently higher than that of `“Random|”`. The minimum GWD of `“Uniform|”` remains unchanged regardless of the number of the trials, which we confirmed by increasing the number of the trials to 10, 000. These results suggest that random initialization provides a more effective optimization pathway, allowing the GWOT to find the better local minima or even global minima.

For both the THINGS and DNN datasets, the minimal GWD and the optimal transportation plan were similar across all the three strategies (Fig. S1a, c and Fig. S2a1-a3, c1-c3), suggesting that these datasets may represent simpler cases where initialization strategies have a minimal impact.

**Fig S1.**
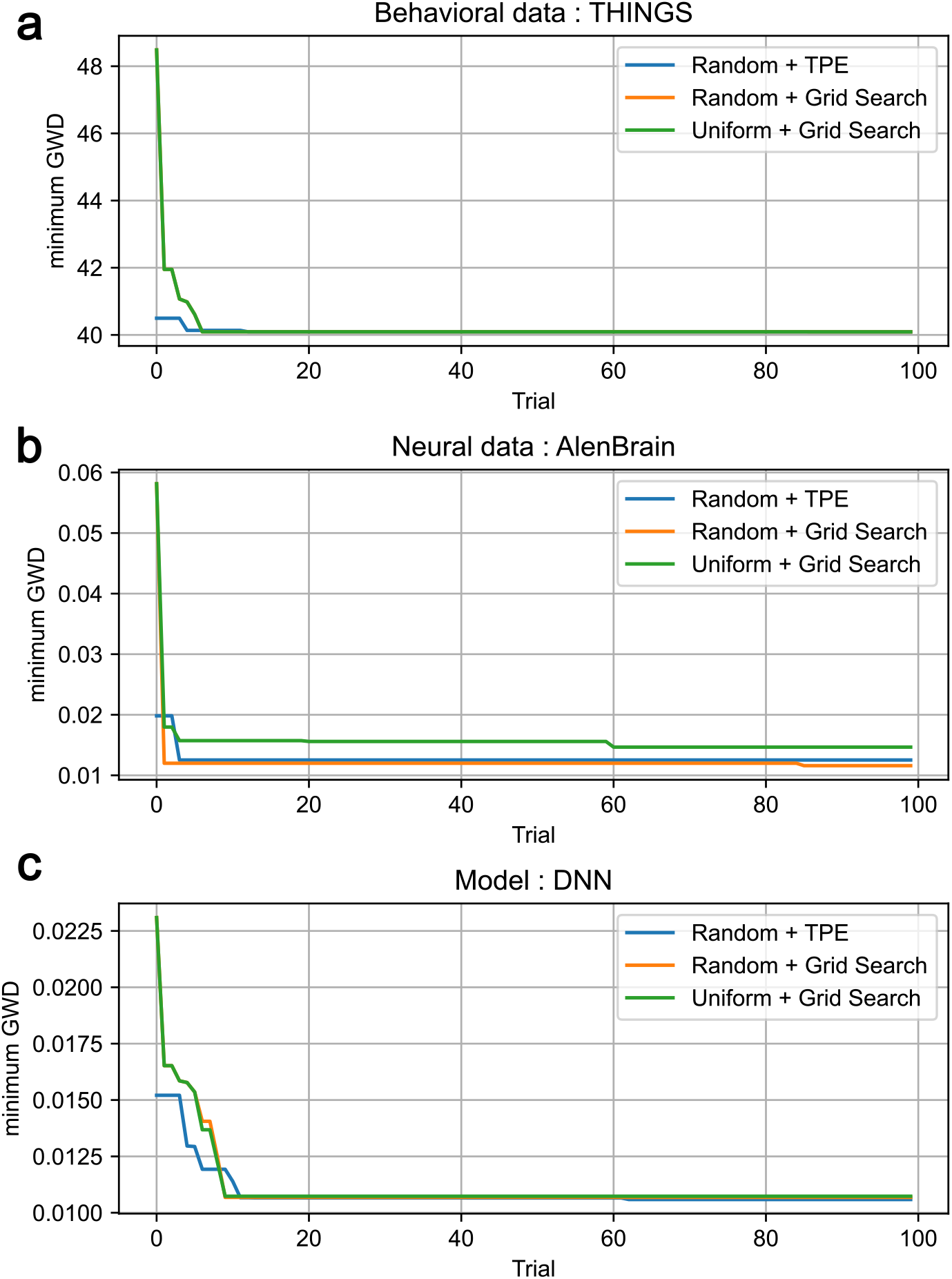
Comparison of the three initialization strategies with real-world data. The change in the minimum GWD value over 100 trials for the three different strategies for each one of three real-world datasets: THINGS, AllenBrain, and DNN. The X-axis shows the number of trials to sample epsilon with its samplers (TPE or Grid Search). The Y-axis shows the minimum GWD value within the trials.

**Fig S2.**
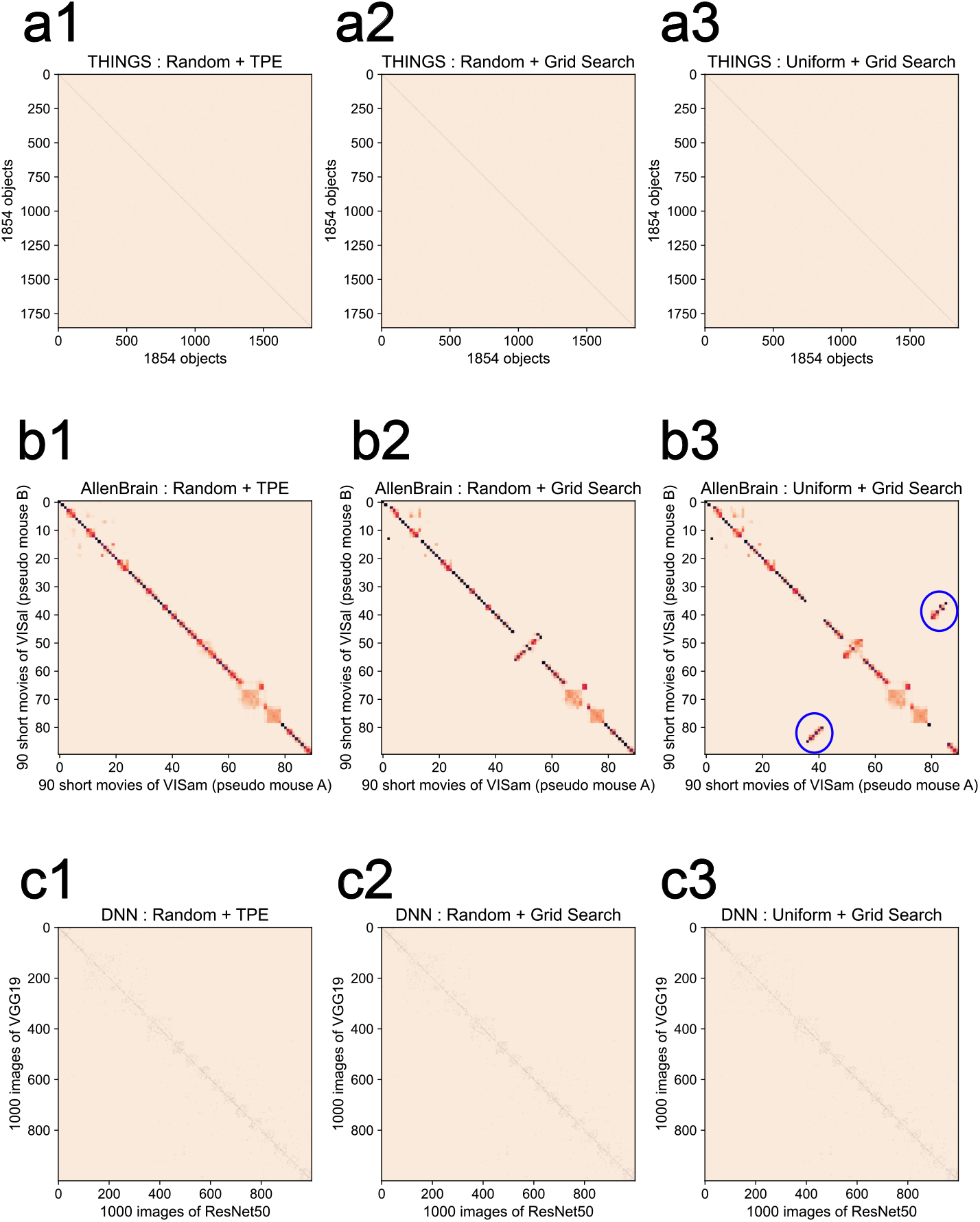
Comparison of optimal transportation plan with minimum GWD obtained through three initialization strategies. The figure a1-a3, b1-b3, and c1-c3 show the optimal transportation plan obtained by the three initialization strategies of each one of three dataset, THINGS, AllenBrain, and DNN, respectively. The blue circles in b3 show some movie chunks are mapped to other chunks in reverse time order (upward to the right).

## References

1. Kriegeskorte N, Mur M, Ruff DA, Kiani R, Bodurka J, Esteky H, et al. Matching categorical object representations in inferior temporal cortex of man and monkey. Neuron. 2008;60(6):1126–1141.

2. Sucholutsky I, Muttenthaler L, Weller A, Peng A, Bobu A, Kim B, et al. Getting aligned on representational alignment. arXiv [q-bioNC]. 2023;2310.13018.

3. Kriegeskorte N, Kievit RA. Representational geometry: integrating cognition, computation, and the brain. Trends Cogn Sci. 2013;17(8):401–412.

4. Haxby JV, Connolly AC, Guntupalli JS. Decoding neural representational spaces using multivariate pattern analysis. Annu Rev Neurosci. 2014;37:435–456.

5. Kawakita G, Zeleznikow-Johnston A, Takeda K, Tsuchiya N, Oizumi M. Is my “red” your “red”?: Unsupervised alignment of qualia structures via optimal transport; 2023. Available from: https://psyarxiv.com/h3pqm/.

6. Mémoli F. Gromov–Wasserstein Distances and the Metric Approach to Object Matching. Found Comut Math. 2011;11(4):417–487.

7. Peyré G, Cuturi M, Solomon J. Gromov-Wasserstein Averaging of Kernel and Distance Matrices. In: Balcan MF, Weinberger KQ, editors. Proceedings of The 33rd International Conference on Machine Learning. vol. 48 of Proceedings of Machine Learning Research. New York, New York, USA: PMLR; 2016. p. 2664–2672.

8. Alvarez-Melis D, Jaakkola TS. Gromov-Wasserstein Alignment of Word Embedding Spaces. arXiv [csCL]. 2018;1809.00013.

9. Alaux J, Grave E, Cuturi M, Joulin A. Unsupervised Hyperalignment for Multilingual Word Embeddings. arXiv [csCL]. 2018;1811.01124.

10. Demetci P, Santorella R, Sandstede B, Noble WS, Singh R. SCOT: Single-Cell Multi-Omics Alignment with Optimal Transport. J Comput Biol. 2022;29(1):3–18.

11. Thual A, Tran H, Zemskova T, Courty N, Flamary R, Dehaene S, et al. Aligning individual brains with Fused Unbalanced Gromov-Wasserstein. arXiv [q-bioNC]. 2022;2206.09398.

12. Kawakita G, Zeleznikow-Johnston A, Tsuchiya N, Oizumi M. Gromov–Wasserstein unsupervised alignment reveals structural correspondences between the color similarity structures of humans and large language models. Sci Rep. 2024;14.

13. Takahashi S, Sasaki M, Takeda K, Oizumi M. Self-supervised learning facilitates neural representation structures that can be unsupervisedly aligned to human behaviors. ICLR 2024 Workshop on Representational Alignment (Re-Align). 2024;.

14. Takeda K, Abe K, Kitazono J, Oizumi M. Unsupervised alignment reveals structural commonalities and differences in neural representations of natural scenes across individuals and brain areas. ICLR 2024 Workshop on Representational Alignment (Re-Align). 2024;.

15. Akiba T, Sano S, Yanase T, Ohta T, Koyama M. Optuna: A Next-generation Hyperparameter Optimization Framework. arXiv [csLG]. 2019;1907.10902.

16. Demetci P, Santorella R, Chakravarthy M, Sandstede B, Singh R. SCOTv2: Single-Cell Multiomic Alignment with Disproportionate Cell-Type Representation. J Comput Biol. 2022;29(11):1213–1228.

17. Hebart MN, Dickter AH, Kidder A, Kwok WY, Corriveau A, Van Wicklin C, et al. THINGS: A database of 1,854 object concepts and more than 26,000 naturalistic object images. PLoS One. 2019;14(10):e0223792.

18. Hebart MN, Zheng CY, Pereira F, Baker CI. Revealing the multidimensional mental representations of natural objects underlying human similarity judgements. Nat Hum Behav. 2020;4(11):1173–1185.

19. Hebart MN, Contier O, Teichmann L, Rockter AH, Zheng CY, Kidder A, et al. THINGS-data, a multimodal collection of large-scale datasets for investigating object representations in human brain and behavior. Elife. 2023;12.

20. Siegle JH, Jia X, Durand S, Gale S, Bennett C, Graddis N, et al. Survey of spiking in the mouse visual system reveals functional hierarchy. Nature. 2021;592(7852):86–92.

21. De Vries SE, Siegle JH, Koch C. Sharing neurophysiology data from the Allen Brain Observatory. Elife. 2023;12:e85550.

22. He K, Zhang X, Ren S, Sun J. Deep residual learning for image recognition. Proceedings of the IEEE conference on computer vision and pattern recognition. 2016; p. 770–778.

23. Simonyan K, Zisserman A. Very Deep Convolutional Networks for Large-Scale Image Recognition. arXiv. 2015;1409.1556.

24. Roads BD, Love BC. Enriching ImageNet with Human Similarity Judgments and Psychological Embeddings. In: Proceedings of the IEEE/CVF Conference on Computer Vision and Pattern Recognition (CVPR); 2021. p. 3547–3557.

25. Gower JC, Dijksterhuis GB. Procrustes Problems. OUP Oxford; 2004.

26. Nili H, Wingfield C, Walther A, Su L, Marslen-Wilson W, Kriegeskorte N. A toolbox for representational similarity analysis. PLoS Comput Biol. 2014;10(4):e1003553.

27. Schütt HH, Kipnis AD, Diedrichsen J, Kriegeskorte N. Statistical inference on representational geometries. Elife. 2023;12.

28. Williams AH, Kunz E, Kornblith S, Linderman S. Generalized shape metrics on neural representations. Advances in Neural Information Processing Systems. 2021;34:4738–4750.

29. Peyré G, Cuturi M. Computational Optimal Transport: With Applications to Data Science. Foundations and Trends® in Machine Learning. 2019;11(5-6):355–607.

30. Flamary R, Courty N, Gramfort A, Alaya MZ, Boisbunon A, Chambon S, et al. POT: Python optimal transport. J Mach Learn Res. 2021;22(1):3571–3578.

31. Chapel L, Alaya MZ. Partial Optimal Tranport with applications on Positive-Unlabeled Learning. Adv Neural Inf Process Syst. 2020;.

32. S’ejourn’e T, Vialard FX, Peyr’e G. The Unbalanced Gromov Wasserstein distance: Conic formulation and relaxation. Adv Neural Inf Process Syst. 2020; p. 8766–8779.

33. Yamins DL, Hong H, Cadieu CF, Solomon EA, Seibert D, DiCarlo JJ. Performance-optimized hierarchical models predict neural responses in higher visual cortex. Proceedings of the national academy of sciences. 2014;111(23):8619–8624.

34. Kell AJE, Yamins DLK, Shook EN, Norman-Haignere SV, McDermott JH. A Task-Optimized Neural Network Replicates Human Auditory Behavior, Predicts Brain Responses, and Reveals a Cortical Processing Hierarchy. Neuron. 2018;98(3):630–644.e16.

35. Zhuang C, Yan S, Nayebi A, Schrimpf M, Frank MC, DiCarlo JJ, et al. Unsupervised neural network models of the ventral visual stream. Proceedings of the National Academy of Sciences. 2021;118(3):e2014196118.

36. Rajalingham R, Issa EB, Bashivan P, Kar K, Schmidt K, DiCarlo JJ. Large-Scale, High-Resolution Comparison of the Core Visual Object Recognition Behavior of Humans, Monkeys, and State-of-the-Art Deep Artificial Neural Networks. J Neurosci. 2018;38(33):7255–7269.

37. Konkle T, Alvarez GA. A self-supervised domain-general learning framework for human ventral stream representation. Nat Commun. 2022;13(1):1–12.

38. Conwell C, Prince JS, Kay KN, Alvarez GA, Konkle T. What can 1.8 billion regressions tell us about the pressures shaping high-level visual representation in brains and machines? bioRxiv. 2023; p. 2022.03.28.485868.

39. Kriegeskorte N, Mur M, Ruff DA, Kiani R, Bodurka J, Esteky H, et al. Matching Categorical Object Representations in Inferior Temporal Cortex of Man and Monkey. Neuron. 2008;60(6):1126–1141.

40. Mur M, Meys M, Bodurka J, Goebel R, Bandettini P, Kriegeskorte N. Human Object-Similarity Judgments Reflect and Transcend the Primate-IT Object Representation. Frontiers in Psychology. 2013;4.

41. Chikazoe J, Lee DH, Kriegeskorte N, Anderson AK. Population coding of affect across stimuli, modalities and individuals. Nat Neurosci. 2014;17(8):1114–1122.

42. Horikawa T, Cowen AS, Keltner D, Kamitani Y. The Neural Representation of Visually Evoked Emotion Is High-Dimensional, Categorical, and Distributed across Transmodal Brain Regions. iScience. 2020;23(5):101060.

43. Koide-Majima N, Nakai T, Nishimoto S. Distinct dimensions of emotion in the human brain and their representation on the cortical surface. Neuroimage. 2020;222:117258.

44. Sagar V, Shanahan LK, Zelano CM, Gottfried JA, Kahnt T. High-precision mapping reveals the structure of odor coding in the human brain. Nat Neurosci. 2023;26(9):1595–1602.

45. Bergstra J, Bardenet R, Kégl B, Bengio Y. Algorithms for Hyper-Parameter Optimization. In: Advances in Neural Information Processing Systems. vol. 24. Curran Associates, Inc.; 2011.

46. Bergstra J, Yamins D, Cox D. Making a Science of Model Search: Hyperparameter Optimization in Hundreds of Dimensions for Vision Architectures. In: Dasgupta S, McAllester D, editors. Proceedings of the 30th International Conference on Machine Learning. vol. 28 of Proceedings of Machine Learning Research. Atlanta, Georgia, USA: PMLR; 2013. p. 115–123.

47. Deng J, Dong W, Socher R, Li LJ, Li K, Fei-Fei L. ImageNet: A large-scale hierarchical image database. In: 2009 IEEE Conference on Computer Vision and Pattern Recognition; 2009. p. 248–255.

